# The aphid BCR4 structure and activity uncover a new defensin peptide superfamily

**DOI:** 10.1101/2021.12.17.472939

**Authors:** Karine Loth, Nicolas Parisot, Françoise Paquet, Catherine Sivignon, Isabelle Rahioui, Mélanie Ribeiro Lopes, Karen Gaget, Gabrielle Duport, Agnès F. Delmas, Vincent Aucagne, Abdelaziz Heddi, Federica Calevro, Pedro da Silva

## Abstract

Aphids (Hemiptera: Aphidoidea) are among the most injuring insects for agricultural plants and their management is a great challenge in agronomical research. A new class of proteins, called Bacteriocyte-specific Cysteine-Rich (BCR), provides an alternative to chemical insecticides for pest control. BCRs have been initially identified in the pea aphid *Acyrthosiphon pisum.* They are small disulfide bond-rich proteins expressed exclusively in aphid bacteriocytes, the insect derived cells that host intracellular symbiotic bacteria. Here, we show that one out of the *A. pisum* BCRs, BCR4, displays an outstanding insecticidal activity against the pea aphid, impairing insect survival and nymphal growth, providing evidence for its potential use as a new biopesticides. Our comparative genomics and phylogenetic analysis indicate that BCRs seem restricted to the aphid lineage. The 3D structure of the BCR4 reveals that this peptide belongs to a yet unknown structural class of peptides and defines a new superfamily of defensins.

## INTRODUCTION

Insects are among the most important pests of cultured plants and stored products, causing an estimated yearly loss of hundreds of millions of dollars worldwide (Oerke & Dehne, 2004; Bradshaw et al., 2016; Diagne et al., 2020). Aphids, in particular, hold a prominent place among insect pests as they represent up to 26% of the pests found on the main food crops grown in temperate climate (maize, wheat, potatoes, sugar beet, barley and tomatoes) (Dedryver et al., 2010; Calevro et al., 2019). Aphid control strategies rely almost exclusively on chemical treatments, which cause persistent environmental pollution and lead to the emergence of insect resistance (Goulson, 2018). New ecologically-friendly solutions are therefore required to control aphids and other phloem-feeders.

Small Disulfide-Rich Proteins (DRP) extracted from plants or arthropods are promising alternative biopesticide molecules (Gressent et al., 2011; Huang et al., 2019; King, 2019). Those naturally occurring molecules display a very broad range of biological activities, mainly related to host-defense processes (Shafee et al., 2017), and show a large structural and chemical diversity. Their exceptional stability is particularly appealing for drug discovery purposes (Gonzalez-Castro et al., 2021). In addition to rather rigid three-dimensional conformations imposed by their polycyclic architecture, DRP exhibit a strong resistance toward *in vivo* enzymatic degradation. Altogether, these features have contributed in establishing DRP as emerging lead compounds for the development of novel peptide-based drugs and, more recently, as potential biopesticides in agronomical research (Rahioui et al., 2014; Huang et al., 2019; King, 2019; Bell et al., 2021). For instance, a knottin DRP extracted from pea seeds, PA1b (Pea Albumin 1, subunit b, 37 amino acids, three disulfide bonds) (Delobel et al., 1998; Gressent et al., 2011; Rahioui et al., 2014), is toxic to numerous insects, including aphids, cereal weevils, mosquitos and moths (Rahioui et al., 2014).

A new class of DRP, called Bacteriocyte-specific Cysteine Rich (BCR) peptides, has recently been identified in a major crop pest, the pea aphid, *Acyrthosiphon pisum* (Shigenobu & Stern, 2013). As many other crop pest insects that thrive on unbalanced diets, aphids have evolved long-lasting relationships with endosymbiotic bacteria and almost all aphids are found in association with the γ3-proteobacterium *Buchnera aphidicola* (Shigenobu et al., 2000; Baumann, 2005; Douglas, 2015) This bacterium supplements the host diet with nutrients lacking or being limited in their habitats, therefore allowing insects to proliferate and cause major economic, social, and health damage. Neither the host nor the endosymbionts can survive independently from each other. *B. aphidicola* is non-culturable and insects artificially deprived of their endosymbionts (aposymbiotic) cannot survive nor reproduce (Brinza et al., 2009). The maintenance of this association relies on the compartimentalization of endosymbionts in specialized insect cells, called bacteriocytes (Simonet et al., 2018). BCRs are encoded by seven orphan genes, and are all expressed exclusively in bacteriocytes of both embryonic and adult aphids (Shigenobu & Stern, 2013). This suggests that BCRs may play a role in bacteriocyte homeostasis, presumably in endosymbiont control as previously in the cereal-weevil endosymbiosis (Login et al. 2011). Consistent with this hypothesis, it has been shown that BCR1, BCR2, BCR3, BCR4, BCR5 and BCR8 exhibit antimicrobial activity or can permeabilize the membrane of *E. coli* cells (Uchi et al., 2019).

Each BCR peptide consists of a secretory signal peptide and a mature peptide, composed of 44-84 amino acids and containing from six in the case of BCR1 to 5 and BCR8, to eight, in the case of BCR6, cysteine residues (Shigenobu & Stern, 2013). Between BCR1, BCR2, BCR4 and BCR5, the cysteine-rich region is highly divergent, but the six cysteines have almost identical spacing in the predicted proteins. Three of these genes, BCR1, BCR4 and BCR5, are found within a genomic region of 20 kbp, suggesting that they may have arisen by a recent tandem gene duplication. No similarity could be found between any of the other BCR family genes (Shigenobu & Stern, 2013). Intriguingly, the pea aphid BCRs show no significant sequence similarity with genes in species outside of the aphid lineage. DRP have undergone extensive divergent evolution in their sequence structure and function. They are classified based on their secondary structure orientation, cysteine distribution across the sequence and cysteine bonding pattern tertiary structure similarities and precursor gene sequence (Shafee et al., 2016; Shafee et al., 2017). Based on their sequence analysis, it was assumed that they could have evolved from defensin-type antimicrobial peptides (AMPs), small proteins containing six to eight cysteines, which are universally found in both animal and plants (Shigenobu & Stern, 2013; Uchi et al., 2019).

In this study, we focused on the BCR4 of *A. pisum*, a benchmark example of peptide with antimicrobial activities (Uchi et al., 2019). We discovered that BCR4 displays an outstanding insecticidal activity against the pea aphid. Tooking advantage of the recent sequencing of several aphid genomes, which enables the study of gene families’ diversification through comparative and evolutionary analyses (Ribeiro Lopes et al., 2020; Huygens et al., 2021), we conducted a comparative genomic analysis across 22 aphid species for which sequence information was available. This allowed the identification of 76 new BCR sequences, all restricted to aphid species and not related to any known defensins. Finally, we determined BCR4 3D structure and showed that it belongs to a new structural class of disulfide-rich proteins. Overall, the biochemical analyses, the evolutionary history and the 3D structure of the BCR4 have highlighted significant insights into the biological and structural properties of BCRs, and have provided evidence for the use of this new defensin superfamily as a potential new biopesticides.

## RESULTS

### Total synthesis of BCR4

To investigate the biological activity of members of the BCR peptide family, a pure sample of synthetic BCR4 was produced through total chemical synthesis. As standard automated Fmoc/*t*Bu solid phase peptide synthesis (SPPS) was unsuccessful (SI appendix Figures S1 and S2, table S1), we turned to a native chemical ligation (NCL)-based approach (Dawson et al., 1994), relying on the assembly of two medium-sized peptide segments. Using a recently developed methodology (Lelievre et al., 2016; Terrier et al., 2016; Terrier et al., 2017; Abboud & Aucagne, 2020; Abboud, et al., 2021a; Abboud, et al., 2021b), we coupled a 20 amino acid (aa) *N*-2-hydroxy-5-nitrobenzylcysteine (*N*-Hnb-Cys) crypto-thioester with a 30 amino acid cysteinyl peptide and obtained the full length reduced form of BCR4 at a high purity (Figure 1). Oxidative folding under thermodynamic control was achieved using a standard protocol (Derache et al., 2012; Martinez et al., 2016), leading to one major compound featuring three disulfide bridges as evidenced by HRMS analysis (SI appendix Figures S3 to S11).

**Figure 1.**
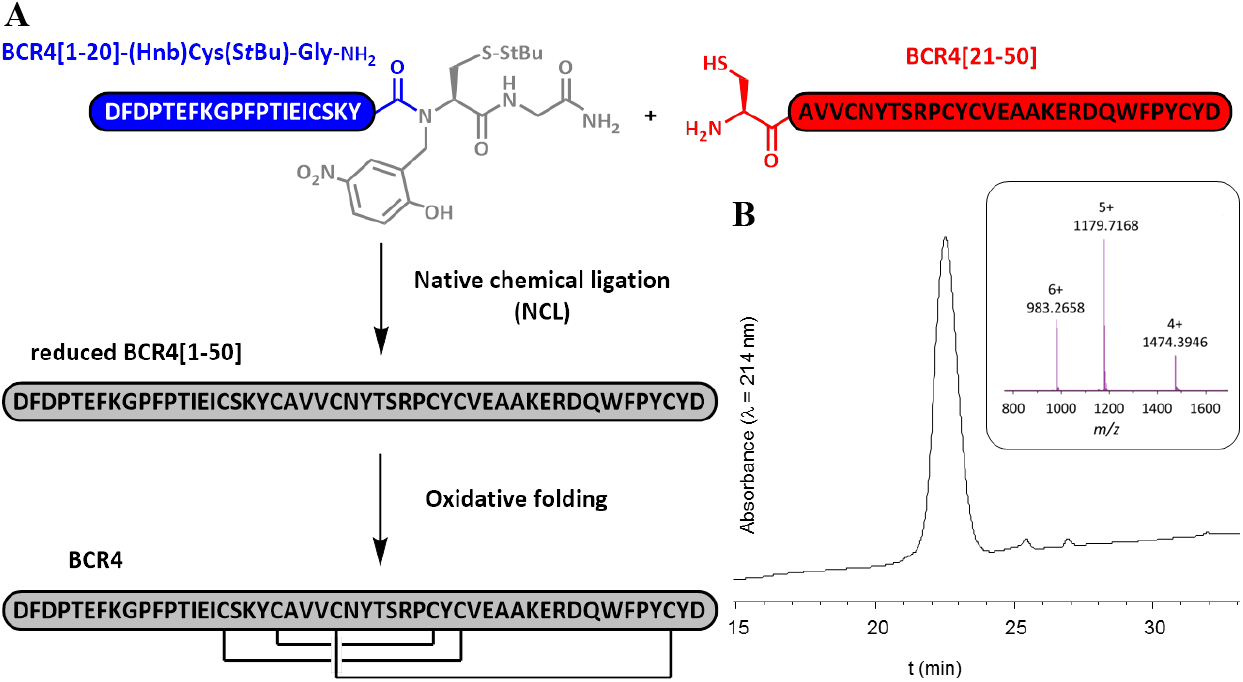
Chemical synthesis of BCR4. **(A)** Schematic representation of the BCR4 peptide chemical synthesis process. **(B)** RP-HPLC chromatogram and ESI-HRMS mass spectrum of the purified folded peptide.

### Antimicrobial activities of BCR4

To assess the antimicrobial activity of BCR4, we tested the effect of various concentrations (ranging from 5 μM to 80 μM) of this peptide on the growth of the Gram-negative bacteria *Escherichia coli* (strain NM522) and the Gram-positive bacteria *Micrococcus luteus.* Consistent with previous observations (Uchi et al., 2019), a slight inhibition of *E.coli* growth was detected at 5 μM of BCR4. This antimicrobial activity increased with BCR4 concentration and we determined a minimal inhibitory concentration (MIC) of 17.0 ± 2.4 μM for which no bacterial growth was detected. Comparatively, no antimicrobial activity was detected against the Gram-positive bacteria *Micrococcus luteus* (Table 1).

**Table 1.** Antimicrobial activities of BCR4 peptide

| | MIC<sup>a</sup> (μM) |
| --- | --- |
| *Escherichia coli* NM522 | $17.0 \pm 2.4$ |
| *Micrococcus luteus* | > 80 |

<sup>a</sup> Minimal inhibitory concentration

### Insect bioassays

The insecticidal potential of BCR4 was assayed by oral administration of various concentration of this peptide to the pea aphid and monitoring of its survival. For all BCR4 concentrations tested (5-80 μM), aphid mortality rate was always significantly greater in the BCR4-aphids compared to the control group, demonstrating a specific effect of BCR4 ingestion on aphid survival (Figure 2). This effect wad dependent on both BCR4 dosage and duration of treatment. The highest concentration tested (80 μM) also had the strongest lethal effect with a Lethal Time 50% (LT50) of 1.16 days (Table 2) and no surviving aphids after two days of BCR4 treatments (Figure 2).

**Figure 2.**
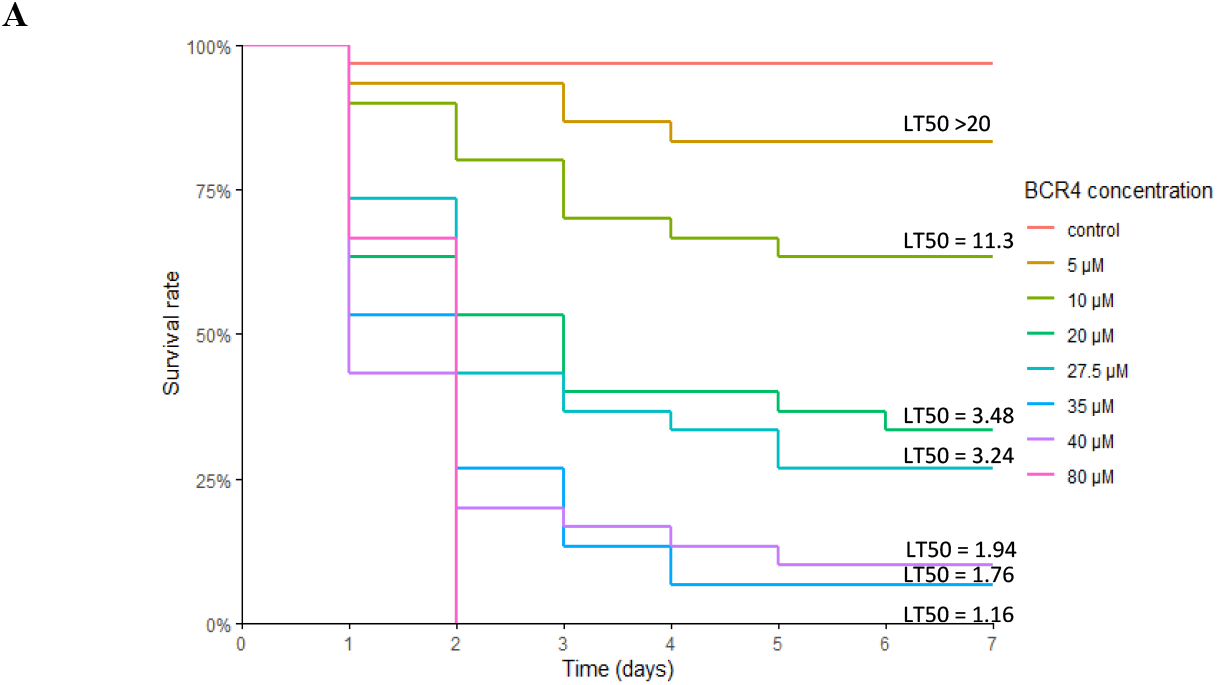
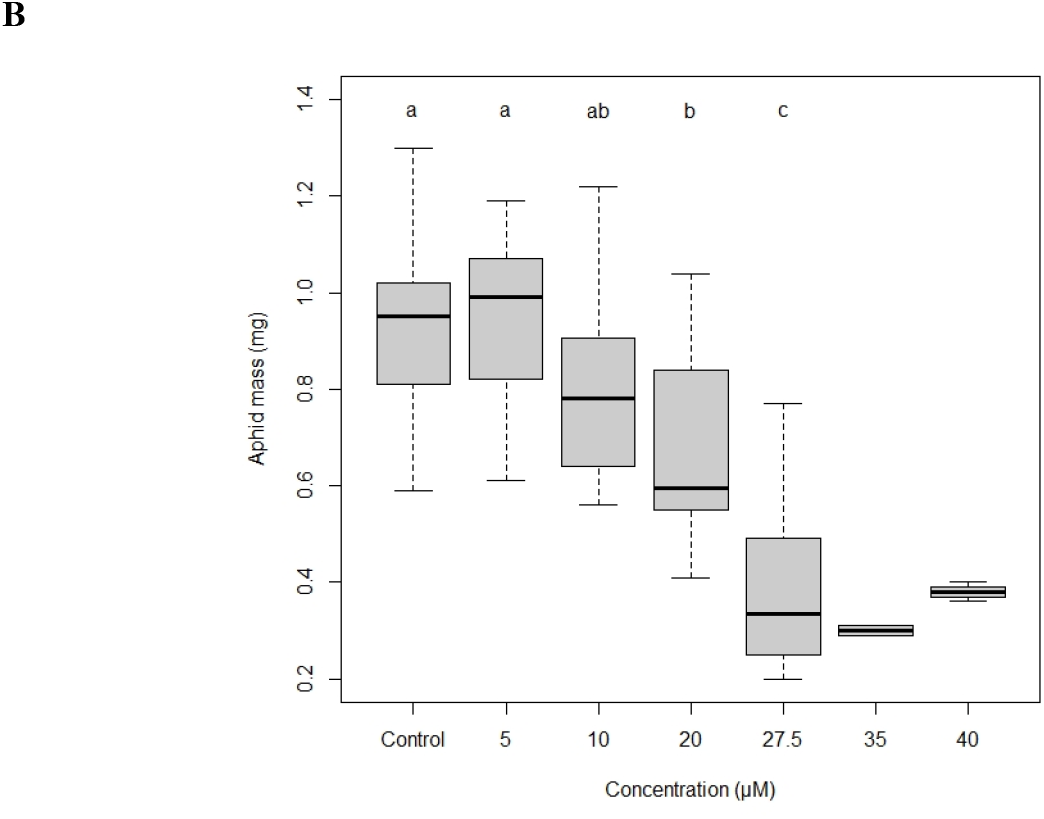
**(A)** Survival curves of the aphid *Acyrthosiphon pisum* reared on artificial diets containing different concentrations of BCR4 peptide. Means Lethal Time 50 (LT50), in days, are indicated above each curve. **(B)** Mass (mg) of 7-day old pea aphid *Acyrthosiphon pisum* subjected to BCR4 treatment. Concentrations labelled with different letters are significantly different (P < 0.05); sample size of BCR4 35 and 40 μM concentrations too small for statistics.

**Table 2.** Toxicity of BCR4 peptide on the pea aphid *Acyrthosiphon pisum* (means Lethal Time 50 (LT50) in days, with confidence intervals), analyzed by survival analysis with a log-normal fit.

|  | Concentration (µM) |  |  |  |  |  |  |
| --- | --- | --- | --- | --- | --- | --- | --- |
| BCR4 peptide | 80 | 40 | 35 | 27.5 | 20 | 10 | 5 |
| LT50 (days) | 1.16<br>[0.98 – 1.37] | 1.76<br>[1.27 – 2.42] | 1.94<br>[1.47 – 2.57] | 3.24<br>[2.22 – 4.72] | 3.48<br>[2.28 – 5.31] | 11.3<br>[4.16 – 30.8] | > 20 |

Even for concentration as low as 20 μM, aphids, on average, did not survive more than 4 days and only one third reached the adulthood. Apart from this effect on aphid survival, the most striking phenotypical effect observed with BCR4 treatment was a statistically significant growth inhibition of the surviving aphids for doses higher than 5 μM, with, for instance, a 60% weight reduction at 7 days after ingestion of BCR4 at 27.5 μM (Figure 2B).

### Phylogeny of the BCR family

To decipher the molecular evolution of the BCR peptides, we aimed at identifying the whole set of proteins homologous to the seven *A. pisum* BCR proteins. Through an exhaustive search of the NCBI nucleotide and protein databases, coupled with a specific inspection of the sequenced aphid species available in the AphidBase (Legeai et al., 2010) database, we retrieved a total of 76 new BCR sequences across 20 aphid species (Table S2) among the 22 for which sequences are available in those publicly available databases. We also found one additional sequence from *A. pisum,* shifting the total number of BCRs in this insect to eight. Interestingly, all these sequences belong to member of the Aphidoidea super-family, thus confirming BCR proteins are restricted to the aphid (*s.l.*) lineage (Shigenobu & Stern, 2013). The complete set of 83 BCR sequences was used for amino acid sequences alignment and phylogenetic tree reconstruction (Figure 3A). Based on these, the BCR sequences can be grouped into four subfamilies including homologs of the BCR1-2-4-5, BCR3, BCR6 and BCR8 sequences of *A. pisum,* respectively. The majority of subfamilies has six cysteine residues with the exception of the BCR6 subfamily which has 8 cysteines. The spacing between cysteines within the BCR sequences is fully conserved within subfamilies, which ensures secure within-family homologies and phylogenetic placement (Table 3).

**Figure 3.**
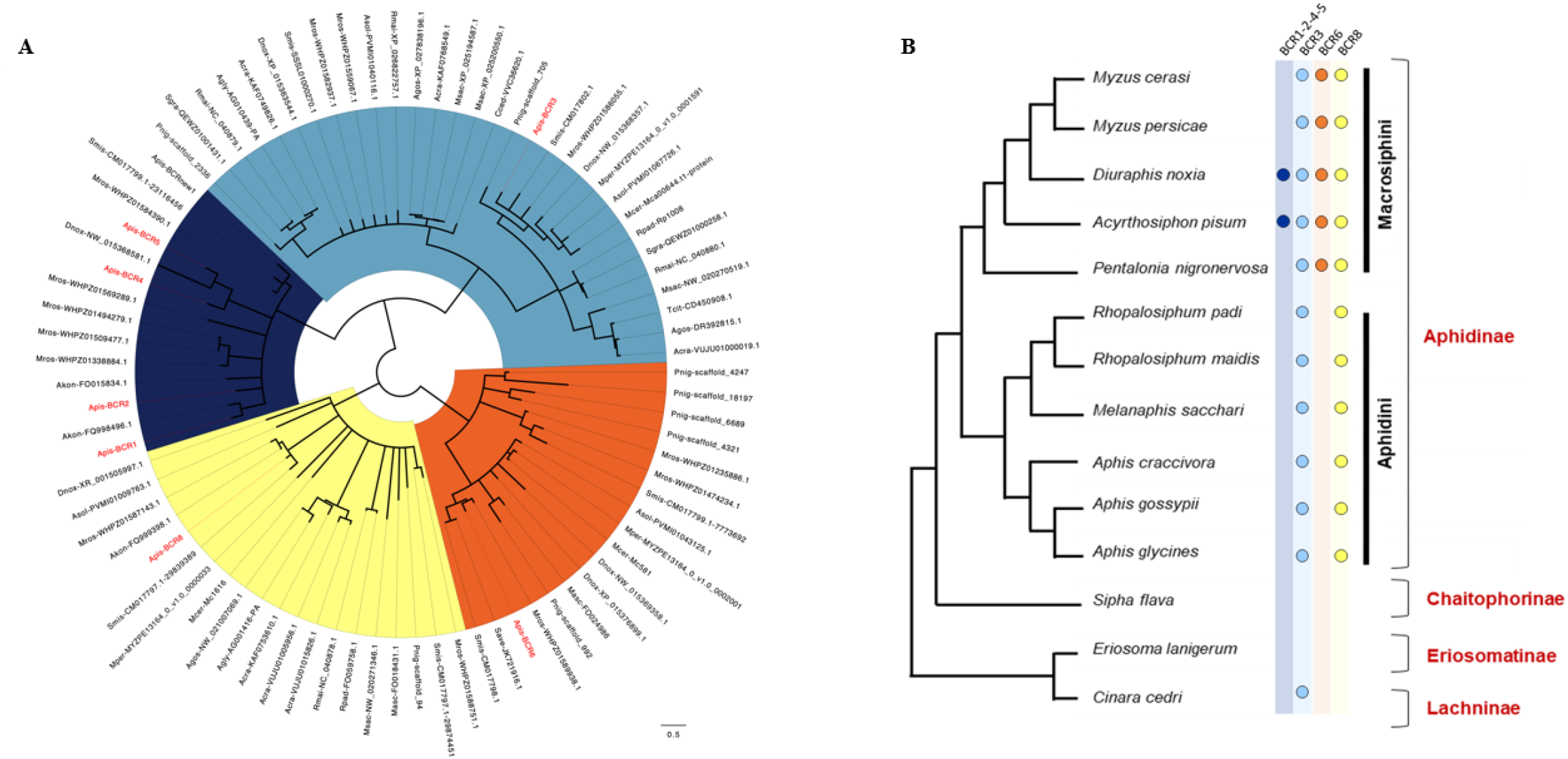
**(A) Phylogenetic trees of BCR proteins**. Maximum-likelihood phylogenetic tree reconstruction was performed using PhyML (Guindon et al., 2010) with an LG 4-rate class model. Branch-support values were calculated using the bootstrap method, with 1000 replicates. Poorly supported branches (<50%) were collapsed using TreeCollapseCL4 (Hodcroft, 2021). The sequences used for the phylogenetic analysis are listed in SI appendix Table S2. Sequences labelled in red reflect those identified by the Shigenobu and Stern (Shigenobu & Stern, 2013) study in the pea aphid *(A. pisum* genome V3.0). Abbreviations: Acra, *Aphis craccivora;* Agly, *Aphis glycines;* Agos, *Aphis gossypii;* Akon, *Acyrthosiphon kondoi;* Apis, *Acyrthosiphon pisum;* Cced, *Cinara cedri;* Dnox, *Diuraphis noxia;* Dvit. *Daktulosphaira vitifoliae;* Masc, *Myzus ascalonicus;* Mcer, *Myzus cerasi;* Mper, *Myzus persicae;* Msac, *Melanaphis sacchari;* Rmai, *Rhopalosiphum maidis;* Rpad, *Rhopalosiphum padi;* Save, *Sitobion avenae;* Sgra, *Schizaphis graminum;* Tcit, *Toxoptera citricida.* **(B) Repartition of BCR peptides in aphid taxonomic groups**. Phylogeny of the 14 members of the aphid group whose genome is annotated and available in databases (among the 19 with sequenced genomes). Their distribution in subfamilies and tribes are indicated in red and black, respectively [adapted from Calevro et al. (2019)]. Colored dots near each species name indicate if a member of each BCR subfamily is present in the genome. Colored dots near each species name indicate if a member of each BCR subfamily is present in the genome of this aphid or not.

**Table 3.**
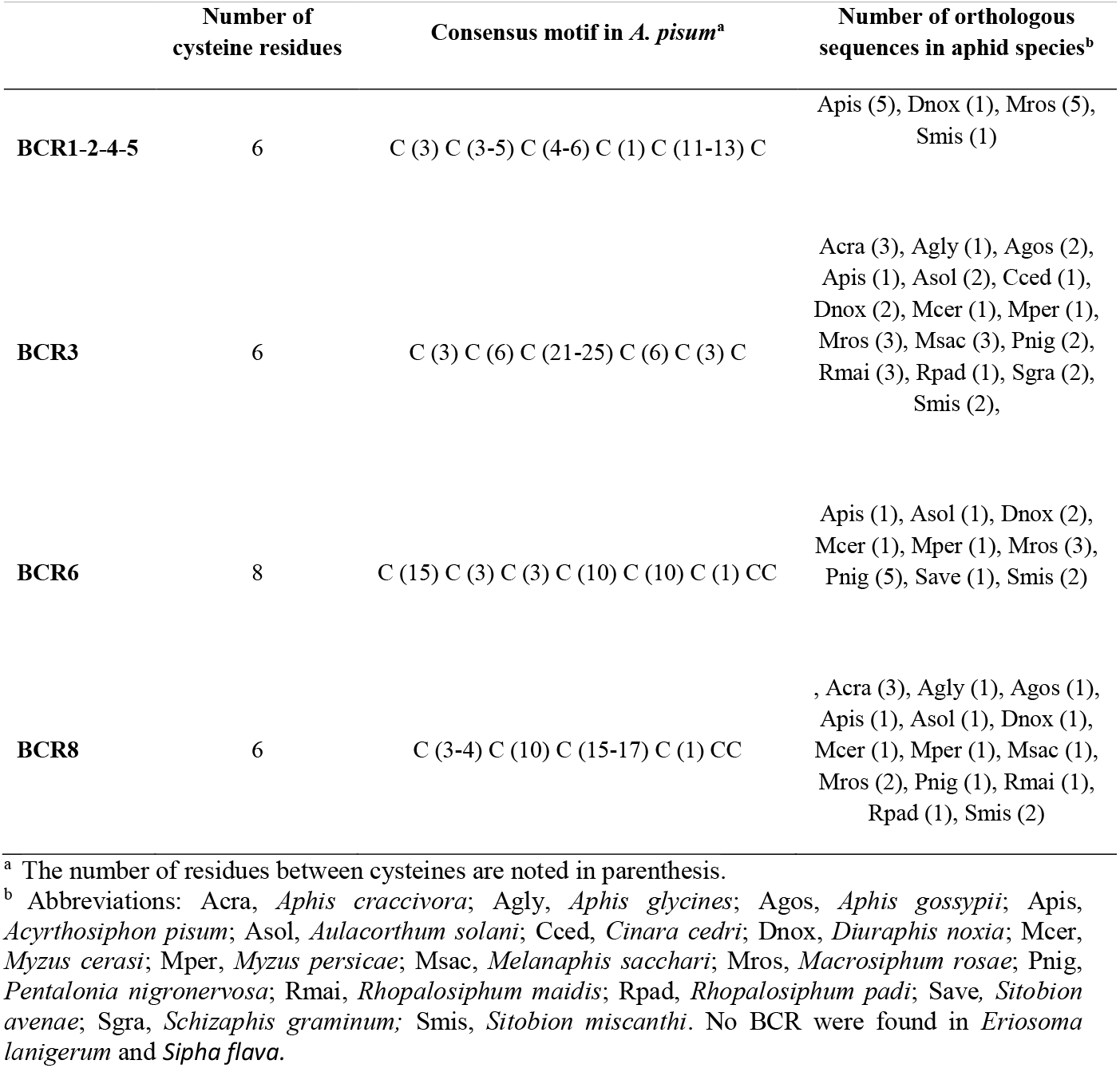
Characteristics of the four BCR subfamilies in 19 aphid species with fully sequenced genomes.

|  | Number of<br>cysteine residues | Consensus motif in *A. pisum*<sup>a</sup> | Number of orthologous<br>sequences in aphid species<sup>b</sup> |
| --- | --- | --- | --- |
| **BCR1-2-4-5** | 6 | C (3) C (3-5) C (4-6) C (1) C (11-13) C | Apis (5), Dnox (1), Mros (5),<br>Smis (1) |
| **BCR3** | 6 | C (3) C (6) C (21-25) C (6) C (3) C | Acra (3), Agly (1), Agos (2),<br>Apis (1), Asol (2), Cced (1),<br>Dnox (2), Mcer (1), Mper (1),<br>Mros (3), Msac (3), Pnig (2),<br>Rmai (3), Rpad (1), Sgra (2),<br>Smis (2), |
| **BCR6** | 8 | C (15) C (3) C (3) C (10) C (10) C (1) CC | Apis (1), Asol (1), Dnox (2),<br>Mcer (1), Mper (1), Mros (3),<br>Pnig (5), Save (1), Smis (2) |
| **BCR8** | 6 | C (3-4) C (10) C (15-17) C (1) CC | , Acra (3), Agly (1), Agos (1),<br>Apis (1), Asol (1), Dnox (1),<br>Mcer (1), Mper (1), Msac (1),<br>Mros (2), Pnig (1), Rmai (1),<br>Rpad (1), Smis (2) |

<sup>a</sup> The number of residues between cysteines are noted in parenthesis.
<sup>b</sup> Abbreviations: Acra, *Aphis craccivora*; Agly, *Aphis glycines*; Agos, *Aphis gossypii*; Apis, *Acyrtosiphon pisum*; Asol, *Aulacorthum solani*; Cced, *Cinara cedri*; Dnox, *Diuraphis noxia*; Mcer, *Myzus cerasi*; Mper, *Myzus persicae*; Msac, *Melanaphis sacchari*; Mros, *Macrosiphum rosae*; Pnig, *Pentalonia nigronervosa*; Rmai, *Rhopalosiphum maidis*; Rpad, *Rhopalosiphum padi*; Save, *Sitobion avenae*; Sgra, *Schizaphis graminum*; Smis, *Sitobion miscanthi*. No BCR were found in *Eriosoma lanigerum* and *Sipha flava*.

Nineteen aphid species have full genome sequences available (*Acyrthosiphon pisum, Aphis craccivora, Aphis glycines, Aphis gossypii, Aulacorthum solani, Cinara cedri, Diuraphis noxia, Eriosoma lanigerum, Macrosiphum rosae, Melanaphis sacchari, Myzus cerasi, Myzus persicae, Pentalonia nigronervosa, Rhopalosiphum maidis, Rhopalosiphum padi, Schizaphis graminum*, *Sipha flava, Sitobion avenae, Sitobion miscanthi*). This permits to predict the complete set of BCR peptides encoded by those aphid genomes. In this group, we observed a high variability in the number and distribution of BCR peptides among the four subfamilies (Table 3). No BCR were found in the *E. lanigerum* (Eriosomatinae subfamily) and *S. flava* (Chaitophorinae subfamily) genomes and only one in *C. cedri* ‘s (Lachninae subfamily) (Figure 3B). Comparatively, all members of the Aphidinae subfamily appear to possess at least two BCR genes and *M. rosae* has the largest repertoire of BCR sequences, with 13 distinct sequences distributed across the four BCR subfamilies. While members of the BCR3 and BCR8 subfamilies are present in all the Aphidinae species included in this study, members of the BCR1-2-4-5 and BCR6 subfamilies appear restricted to the Macrosiphini aphid tribe suggesting that the genes encoding those BCRs arose from duplication of the former.

### BCR4 solution structure

To better characterise the structure evolution of the BCR4 peptide, we have determined its 3D structure by NMR spectroscopy. The ^1^H NMR and the natural-abundance ^1^H-^15^N sofast-HMQC spectra (Schanda et al., 2005) of the protein showed a good dispersion of the amide chemical shifts, indicative of highly structured peptides. Following standard procedures, the analysis of the set of 2D-TOCSY and NOESY spectra allowed a complete assignment of ^1^H chemical shifts. This assignment was facilitated by heteronuclear ^1^H-^15^N and ^1^H-^13^C NMR spectra, particularly in crowded regions of the ^1^H TOCSY and NOESY spectra corresponding to side chains (BRMB entry 34197). The 3D structures were calculated by considering a total of 923 distance restraints, 16 hydrogen bonds, 88 dihedral angles and 3 ambiguous disulfide bridges (Table 4).

**Table 4.** NMR constraints and structural statistics.

| NMR restraints |  |
| --- | --- |
| *Distance restraints* |  |
| Total NOE | 923 |
| Unambiguous | 763 |
| Ambiguous | 160 |
| Hydrogen bonds | 16 |
| *Dihedral Angle Restraints* | 88 |
| *Disulfide bridges<sup>a</sup>* | 3 |
| Structural Statistics (7PQW.pdb) |  |
| *Average violations per structure* |  |
| NOEs $\geq 0.3 \text{ \AA}$ | 0 |
| Hydrogen bonds $\geq 0.3 \text{ \AA}$ | 0 |
| Dihedrals $\geq 15^\circ$ | 3 |
| Dihedrals $\geq 10^\circ$ | 8 |
| Average pairwise rmsd ($\text{\AA}$) | 0.206 |
| Ramachandran Analysis |  |
| Most favored region and allowed region | 96.9 |
| Generously allowed | 0.7 |
| Disallowed | 2.4 |
| Energies ($\text{kcal.mol}^{-1}$)<sup>b</sup> | |
| Electrostatic | $-1631.48 \pm 29.53$ |
| Van der Waals | $-334.18 \pm 11.14$ |
| Total energy | $-1196.56 \pm 31.80$ |
| Residual NOE energy | $56.38 \pm 5.28$ |

<sup>a</sup> Introduced as ambiguous; <sup>b</sup> Values are given as mean ± standard deviation (n=20)

The 200 water-refined structures of BCR4 possess three disulfide bridges with an identical pairing: C17-C34, C21-C32 and C25-C48. Among them, 20 structures were selected in agreement with all NMR experimental data and the standard covalent geometry. Coordinates were deposited as PDB entry 7PQW. Analysis of the 20 final structures with PROCHECK-NMR (Laskowski et al., 1996) showed that almost all of the residues (96.9%) were in the most favored or additionally allowed regions of the Ramachandran diagram (Table 4).

BCR4 is folded into a compact globular unit consisting of an N-terminal tail (D1-T13) and an α-helix (T14-V24) followed by an antiparallel β-sheet composed of two short β-strands, β_1_ C34-A37 and β_2_ Q43-P46 (Figure 4). The 3D structure is stabilized by three disulfide bridges linking the α-helix to β_1_ (C17-C34), to the loop C25-Y33 (C21-C32) and to the C-terminal tail (C25-C48), respectively.

**Figure 4.**
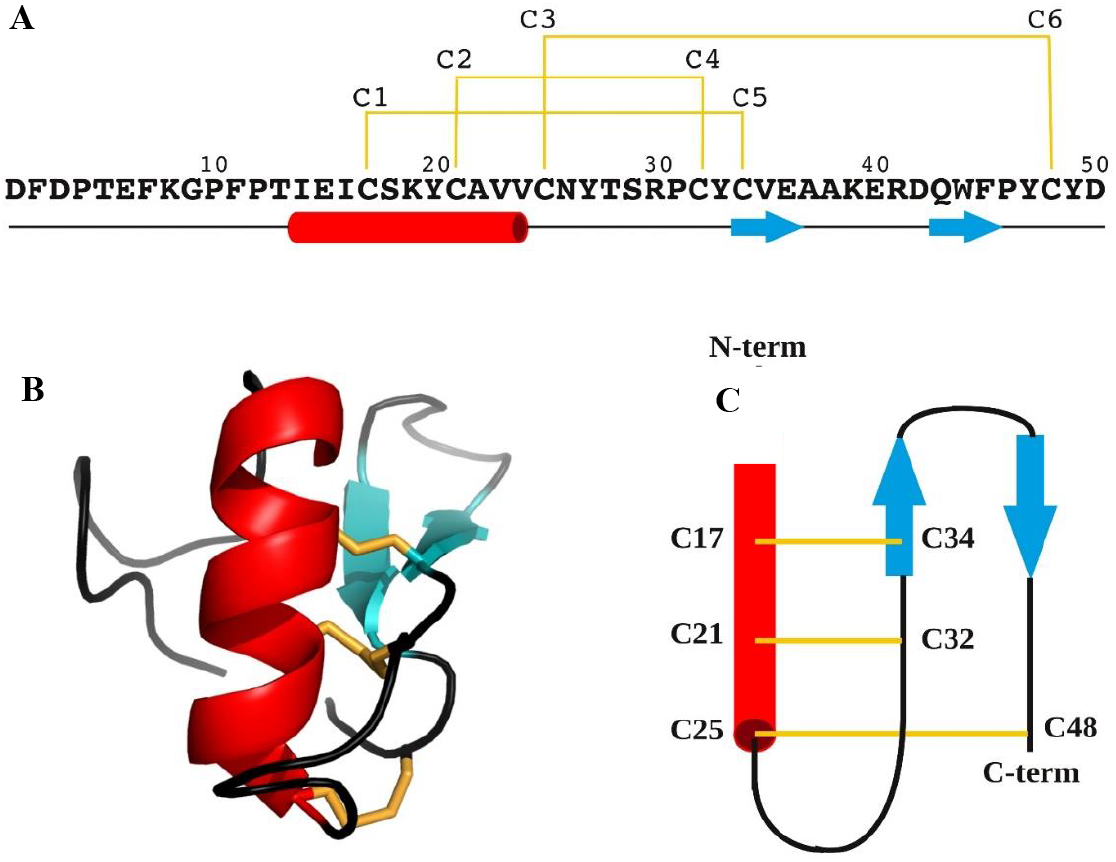
**(A)** Primary structure of BCR4 peptide. **(B)** Three-dimensional structure of BCR4. **(C)** Topology diagram of BCR4. α-helix, β-sheet, and random coil are represented in red, cyan and black, respectively. Disulfide bonds are colored in yellow.

## DISCUSSION

Thus far, most pest control strategies have relied on the use of systemic chemical pesticides, which are increasingly stigmatized because of their persistence in the environment and their toxicity to non-target organisms (Goulson, 2018). This creates the need for the development of new pest management solutions, which will undoubtedly be built upon a better investigation of new biological molecules that may be used as potential biopesticides. Small Disulfide Rich Proteins (DRPs) extracted from plants or arthropods are promising alternative biopesticide molecules (Rahioui et al., 2014; Huang et al., 2019; King, 2019). Genomic plasticity of DRP-encoding sequences is known to foster the adaptability of organisms and to enable the acquisition of new functions (Shafee et al., 2017). In this work, we successfully produced in sufficient amount and purity the folded form of BCR4 (Figure 1), a DRP encoded by the pea aphid genome and part of the BCR family of peptides, that was used to explore the insecticidal properties of this protein. We showed that BCR4 strongly interferes with survival and growth of *A. pisum* in a dose-dependent manner (Figure 2). Its range of activity (5-80 μM, Figure 2) is similar to that displayed by PA1b, a promising plant biopesticide (Gressent et al., 2011; Rahioui et al., 2014) active on the same insect target (Gressent et al., 2007). Importantly, from a functional perspective and as previously reported (Uchi et al., 2019), BCR4 has a significant bactericidal effect on *E. coli* (Table 1), a free-living relative of *B. aphidicola*, the obligatory aphid endosymbiont (Uchi et al., 2019). Based on these results, we propose that the ingestion of exogenous BCR may block nymphal development and induce aphid death by interfering with the population density of this endosymbiont, essential for nymphal growth and survival (Brinza et al., 2009).

From an evolution perspective, a large-scale homology searches through genomic, transcriptomic and proteomic databases was performed to complete the repertoire of BCR peptides, up to now limited to seven identified sequences in the pea aphid genome. Importantly, the 76 additional sequences we found (bringing to 83 the total number of BCR sequences identified) are all encoded by aphid genomes (Figure 3). Their evolutionary analysis showed that aphid BCRs are organized into four subfamilies including BCR1-2-4-5, BCR3, BCR6 and BCR8 sequences (Figure 3), respectively. BCRs from those four subfamilies all bear 6 or 8 cysteine residues and we observed a very clear intra-group conservation of cysteine topology (e. g. distribution in the sequence and disulfide pairing). But, as for other DRP, BCRs present high sequence diversity in their inter-cysteine loops preventing the detection of any firm homology with arthropod defensins at this moment (Shafee et al., 2016; Shafee et al., 2017).While BCR subfamilies had been previously reported (Shigenobu & Stern, 2013), we were able to enrich each of them with many new members (Figure 3). We also showed that (i) aphids present varying numbers of BCR-encoding genes, from 1 to 13 and (ii) while BCR3 and BCR8 homologs are wildly spread in the aphid lineage, genes from the BCR1-2-4-5 and BCR6 subfamilies are restricted to the Macrosiphini tribe, which includes many major agricultural pests. This suggests a complex evolutionary history involving several events of gene duplications and losses with possible functional diversification of the resulting homologs.

To study the structure evolution of the BCR4 peptide, its three-dimensional structure was determined by NMR spectroscopy and careful protein fold analysis revealed that BCR4 belongs to a yet unknown structural class of defensin proteins (Figure 4). Defensins are a well-characterized group of DRPs present in all eukaryotes’ genomes (Shafee et al., 2016). Two superfamilies have been described to date, each derived from independent evolutionary event: the so-called trans-defensin superfamily is uniquely composed of the CSαβ (cysteine-stabilized α-helix β-sheet) family and the cis-defensin superfamily is composed of the α-, β-, θ-, and big defensins families (Shafee et al., 2016; Shafee et al., 2017). Contrary to the CSαβ family (Figure 5), the CXC motif (X denoting any amino acid residue) of BCR4 is in the first β-strand, leading to a different cysteine bonding pattern, the one corresponding to the β-defensin fold (the second family of defensin well-known as antimicrobial peptides) (Shafee et al., 2016; Shafee et al., 2017). The fold of BCR4 is therefore a new type of defensin peptide suggesting that BCRs constitute a new class of antimicrobial cysteine-rich proteins. Shaffee’s phylogenetic work showed that the defensins consist of two independent and convergent superfamilies (Shafee et al., 2016; Shafee et al., 2017). We here hypothesize that the defensins consist of at least three independent and convergent superfamilies: cis-, trans- and BCR defensins

**Figure 5.**
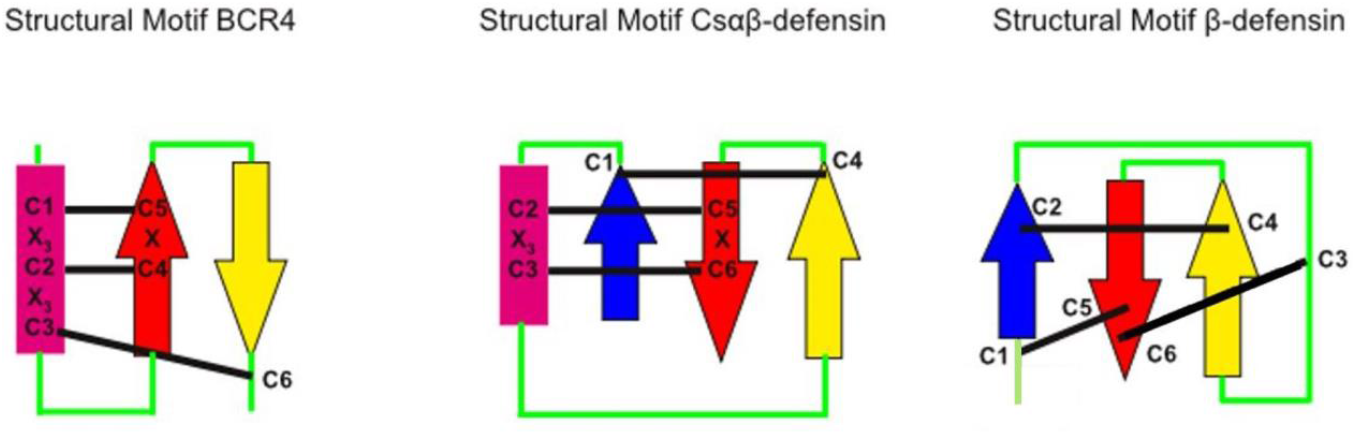
Disulfide connectivities in BCR4 peptide and in CSαβ and β defensin groups. α-helices indicated by rectangle, β-strands represented by arrows, and disulfides in black lines.

BCRs are orphan genes exclusively expressed in aphid bacteriocytes. This gene family has probably evolved in aphid lineages to ensure several functions related to endosymbiosis, including bacteriocyte homeostasis and endosymbiont control. Related studies on cereal weevils *Sitophilus* have shown that the coleoptericin A (ColA) AMP selectively targets endosymbionts within bacteriocytes and impairs bacterial cytokinesis, thereby regulating bacterial cell division and preventing bacterial exit from weevil bacteriocytes (Login et al., 2011; Login & Heddi, 2013). Similar results were obtained in the *Rhizobium*-legames symbiosis, where nodule-specific secreted peptides called NCRs were shown to target bacteria and induce an irreversible elongation of bacteria, rendering them metabolically active but unable to multiply *in vitro* outside plant nodules (Van de Velde et al., 2010). In the actinorhizal plant *Alnus glutinosa*, nodules express defensin-like peptides, including the Ag5 peptide that targets symbiont vesicles and increases permeability of vesicle membranes (Carro et al., 2015). Small peptides are becoming emerging molecules that target a variety of functions in host-symbiont interaction, including symbiont control and physiology (Mergaert et al., 2017). Such a convergent evolution pattern is most interesting in its applied perspectives.

Overall, the susceptibility of aphids to BCR peptides may lead to the development of effective strategies for controlling such sucking pests. The exploration of this work may end up in a new protein family lead for the specific control of aphids, which include some of the most important pests of global agriculture.

## Materials and Methods

### Peptide synthesis

BCR4 was synthesized through native chemical ligation (NCL) of two peptide segments, followed by oxidative folding to form the three disulfide bridges, using already described protocols (Lelievre et al., 2016; Terrier et al., 2016; Terrier et al., 2017; Abboud & Aucagne, 2020; Abboud et al., 2021a; Abboud et al., 2021b). Briefly, a N-terminal cysteinyl peptide segment (sequence: H-^21^CAVVCNYTSRPCYCVEAAKERDQWFPYCY^50^D-OH) and a C-terminal crypto-thioester segment (sequence: H-^1^DFDPTEFKGPFPTIEICSK^20^Y- (Hnb)C(StBu)G-NH2) were synthesized on a Prelude peptide synthesizer (Protein Technologies, Tucson, AZ, USA) using standard Fmoc/tBu chemistry at a 25 μmol scale and starting from a Tentagel resin equipped with a Rink’s amide linker. The *N*-2-lιvdiOw-5-nitrobenzyl (Hnb) group was automatically introduced through resin reductive amination. Both peptide segments were purified by C_18_ reverse phase (RP)-HPLC on a Nucleosil C_18_ column (300 Å, 250 × 10 mm) (Macherey-Nagel, Düren, Germany), maintained at 25°C. Solvent A was 0.1% trifluoroacetic acid (TFA) in water and B was 0.1% TFA in acetonitrile. A linear gradient, from 20% B to 55% B, was applied for 30 min at a flow rate of 3 ml/min and afforded the pure reduced form of BCR4 through collection of the peak identified as the target compound. The purified peptides were then chemoselectively coupled under standard NCL conditions (2 mM peptide segments, 50 mM tris-carboxyethylphosphine, 200 mM 4-mercaptophenylacetic acid, 6 M guanidinium chloride, 200 mM phosphate buffer, pH 6.5, 37 °C, 24 h) (Lelievre et al., 2016; Terrier et al., 2016) and purified as previously to retrieve the pure reduced form of BCR4 (sequence: H-^1^DFDPTEFKGPFPTIEICSKYCAVVCNYTSRPCYCVEAAKERDQWFPYCY^50^D-OH).

The purified compound was further analyzed by ESI-HRMS on a Bruker maXisTM 22 ultra-high-resolution Q-TOF mass spectrometer (Bruker Daltonics, Bremen, Germany), in positive mode and a [M+H]^+^ *m/z* ratio of 5897.5883 was found (theoretical monoisotopic *m/z* calculated for C_266_H_379_N_62_O7_9_S_6_: 5897.5869). Subsequently, the pure reduced form of BCR4 was oxidatively folded *in vitro* (10 μM peptide concentration, 0.1 mM oxidized glutathione, 1 mM glutathione, 1 mM EDTA, 100 mM TRIS, pH 8.5, 20°C, 48 h) (Silva et al., 2009; Derache et al., 2012) before purification to homogeneity by C_18_ RP-HPLC. Analysis of the purified folded product *via* ESI-HRMS analysis gave results consistent with the formation of three disulfide bridges ([M+H]^+^ *m/z* obtained: 5891.5437; theoretical monoisotopic *m/z* calculated for C_266_H_373_N_62_O_79_S_6_: 5891.5400).

### Antimicrobial assays

*E. coli* and *M. luteus* were grown in Lysogeny Broth (LB) and Terrific Broth (TB) medium (Sigma-Aldrich, Saint-Louis, MO, USA), respectively. Antimicrobial assays were performed in sterile 96-well plates with a final volume of 100 μl per well, composed of 50 μl of culture and 50μl of serially diluted peptides (5-80 μM). *E. coli* and *M. luteus* were added at an OD600 of 0.006 and 0.05, respectively. Plates were incubated at 30 °C for 24 h and growth was measured at 600 nm using a Power wave XS-Biotek plate reader (Bioteck Instrument, Colmar, France). The lowest concentration of BCR4 peptide showing complete inhibition was taken as minimal inhibitory concentration (MIC). All analyses were performed in triplicate, with the results expressed as mean±standard deviation of mean (SEM).

### Insects and insect assays

The aphid clone used was *A. pisum* LL01, a long-established alfalfa-collected clone containing only the primary endosymbiont *B. aphidicola.* Aphids were maintained on young broad bean plants (*Vicia faba* L. cv. Aguadulce) at 21 °C, with a photoperiod of 16 h light - 8 h dark to obtain strictly parthenogenetic aphid matrilines, that were reared and synchronized as previously described (Simonet et al., 2018).

For toxicity analyses, three groups of 10 1^st^ instar nymphs (aged between 0 and 24 hours) were collected and placed in *ad hoc* feeding chambers containing an AP3 artificial diet (Febvay et al., 1988) supplemented with different experimental doses (5 μM to 80 μM) of solubilised BCR4 peptide. Toxicity was evaluated by scoring survival daily over the whole nymphal life of the pea aphid (7 days) (Simonet et al., 2016). Growth was also measured by weighing adult aphids on a Mettler AE163 analytical microbalance (Mettler Toledo, Columbus, OH, USA) at the closest 10 μg, following the protocol described previously (Rahbe & Febvay, 1993; Loth et al., 2015).

Aphid mortality data for all BCR4 concentrations were analysed separately in a parametric survival analysis with a log-normal fit. Aphid weights were analysed by Anova followed by Tukey-Kramer HSD test for comparing multiple means. All analyses were made with JMP software version 11 (SAS Institute Cary USA, MacOS version).

### Sequence analysis and phylogeny

Homologous BCR protein sequences were retrieved using a combination of TBLASTN and BLASTP (Altschul et al., 1997; Boratyn et al., 2012) against the aphid genomes available from (i) AphidBase (Legeai et al., 2010), (ii) the NCBI genome database and (iii) the whole NCBI non-redundant protein, nucleotide and EST databases (with *A. pisum* BCR as blast seeds; see SI appendix Table S2 for a complete list of BCR protein sequences used for the phylogenetic analysis). BCR protein sequences were subjected to multiple sequence alignments using the MUSCLE program (Edgar, 2004). Subsequently, a phylogenetic tree was constructed, using the PhyML method (Guindon et al., 2010) implemented in the Seaview software (v5.0.4) (Gouy et al., 2010; Guindon et al., 2010) (LG model with 4 rate classes), and the reliability of each branch was evaluated using the bootstrap method, with 1000 replicates. Poorly supported branches (<50%) were collapsed using TreeCollapseCL4 (Hodcroft, 2021). Graphical representation and editing of the phylogenetic tree were performed with FigTree (v1.4.3) (Drummond & Rambaut, 2007).

### NMR Experiments

Prior to NMR analysis, the synthesized BCR4 peptides were dissolved in H2O:D2O (9:1 ratio) at a concentration of 0.6 mM and pH was adjusted to 4.8. Then, 2D ^1^H NOESY, 2D ^1^H TOCSY, a sofast-HMQC (Schanda et al., 2005) (^15^N natural abundance) and a ^13^C-HSQC (^13^C natural abundance) were performed at 298 K on an Avance III HD BRUKER 700 MHz spectrometer equipped with a cryoprobe. ^1^H chemical shifts were referenced to the water signal (4.77 ppm at 298 K). NMR data were processed using the Topspin software version 3.2^TM^ (Bruker, Billerica, MA, USA) and analyzed with CCPNMR version 2.2.2 (Vranken et al., 2005).

### Structure calculations

Structures were calculated using the Cristallography and NMR System (Brunger et al., 1998; Brunger, 2007) through the automatic assignment software ARIA2 version 2.3 (Rieping et al., 2007) with NOE derived distances, hydrogen bonds (in accordance with the observation of typical long or medium distance NOE cross peaks network for β-sheets and α-helices respectively – HN/HN, HN/Hα, Hα/Hα), backbone dihedral angle restraints (determined with the DANGLE program (Cheung et al., 2010) and three ambiguous disulfide bridges. The ARIA2 protocol, with default parameters used, simulated annealing with torsion angle and Cartesian space dynamics. The iterative process was repeated until the assignment of the NOE cross peaks was complete. The last run for BCR4 was performed with 1000 initial structures and 200 structures were refined in water. 20 structures were selected on the basis of total energies and restraint violation statistics, to represent the structure of BCR4 in solution. The quality of final structures was evaluated using PROCHECK-NMR (Laskowski et al., 1996) and PROMOTIF (Hutchinson & Thornton, 1996). The figures were prepared with PYMOL (De Lano, 2002).

## ACKNOWLEDGEMENTS

This work was supported by INRAE (Institut National de Recherche pour l’Agriculture, l’Alimentation et l’Environnement), INSA Lyon (Institut National des Sciences Appliquées Lyon) and the French ANR-119-CE11-0004-01 (Biofamily) program grant. We thank Philippe Marceau and Jean-Baptiste Madinier for assistance in solid phase peptide synthesis.

We declare no competing interests.

## AUTHORS’ CONTRIBUTIONS

Karine Loth: investigation, formal analysis, data curation, validation, visualisation; Nicolas Parisot: investigation, formal analysis, data curation, validation, visualisation; Françoise Paquet: investigation, formal analysis, data curation, validation, visualisation, Catherine Sivignon: investigation; Isabelle Rahioui : investigation; Mélanie Ribeiro Lopes: formal analysis, visualization; writing—review & editing; Karen Gaget: resources; Gabrielle Duport : resources, Agnès F. Delmas : conceptualisation, writing—review & editing, Vincent Aucagne: conceptualisation, investigation, formal analysis, data curation, validation, visualisation, writing—review & editing, Abdelaziz Heddi writing—review & editing, Federica Calevro conceptualisation, supervision, writing—review & editing, funding Acquisition and Pedro da Silva : conceptualisation, supervision, project administration, writing—original draft, Funding Acquisition

