## Supplemental_file for "The aphid BCR4 structure and activity uncover a new defensin peptide superfamily"

[3]: Univ Lyon, INSA Lyon, INRAE, BF2I, UMR 203, F-69621, Villeurbanne, France

4: Univ Lyon, INRAE, INSA Lyon, BF2I, UMR 203, F-69621, Villeurbanne, France

### These authors contributed equally

*To whom correspondence should be addressed: Pedro Da Silva,

**This PDF file includes:**

Supplementary methods for BCR4 chemical synthesis (including table S1 and Figures S1 to S10)

Table S2

**BCR4 chemical Synthesis**

#### 1. General information

All reagents and solvents were used without further purification. Protected amino acids, Fmoc-Rink amide linker, Fmoc-Tyr(*t*Bu)-Thr($\Psi^{(Me,Me)Pro}$)-OH and HCTU were purchased from Merck Biosciences (Nottingham, UK). Tentagel R $NH_2$ and Wang-type Fmoc-Asp(O*t*Bu) TentaGel R PHB resins were purchased from Rapp polymers (Tuebingen, Germany). Peptide synthesis grade DMF was purchased from VWR (Fontenay-sous-Bois, France). Ultrapure water was obtained using a Milli-Q water system from Millipore (Molsheim, France). All other chemicals were from Sigma Aldrich (St-Quentin-Fallavier, France) and solvents from SDS-Carlo Erba (Val de Reuil, France).

High resolution ESI-MS analyses were performed on a maXis ultra-high-resolution Q-TOF mass spectrometer (Bruker Daltonics, Bremen, Germany), using the positive mode. The multiply-charged envelope was deconvoluted using the Charge Deconvolution algorithm in Bruker Data Analysis 4.1 software to obtain the monoisotopic $[M+H]^+$ molecular ion value. HPLC analyses were carried out on a LaChrom Elite system consisting of an L-2130 pump, an L-2455 diode array detector and an L-2200 autosampler, and equipped with a Jupiter C4 column (300 Å, 5 µm, 250 × 4.6 mm, 1 mL/min flow rate). Semi-preparative HPLC purifications were carried out on a Chromaster 600 system consisting of a 5160 pump, a 5430 diode array detector and a 5260 autosampler, equipped with either a Jupiter C4 (300 Å, 5 µm, 250 × 10 mm, 3 mL/min flow rate) or a Nucleosil C18 (300 Å, 5 µm, 250 × 10 mm, 3 mL/min flow rate) column. Solvents A and B are 0.1% TFA in $H_2O$ and 0.1% TFA in MeCN, respectively. Chromatography was conducted at room temperature unless otherwise mentioned. LC/HRMS analyses were carried out on an Ultimate 3000 RSLC HPLC system (Dionex, Germering, Germany), coupled with the maXis mass spectrometer and fitted with a Aeris WidePore XB-

C18 (200 Å, 3.6 µm, 2.1 × 150 mm, 0.5 mL/min flow rate, 40°C) column. Solvents A and B were 0.1% formic acid in $H_2O$ and 0.08% formic acid in MeCN, respectively. Gradient: 3% B for 0.6 min, then 3 to 50% B over 10.8 min.

Unless specified otherwise, quantities of purified peptides were determined by weight, based on a molecular mass taking into account trifluoroacetate counter-ions (one per Arg, His, Lys and N-terminal amine of the peptide sequence) but not water content.

Deoxygenation of solutions used for native chemical ligation and oxidative folding was performed through four consecutive vacuum (~5 mbar)/argon cycles.

#### 2. General procedures for solid phase peptide synthesis

Fmoc-based solid phase peptide syntheses (SPPS) were carried out on a Prelude synthesizer from Protein Technologies (Tucson, Arizona USA). Standard side-chain protecting groups were used: Arg(Pbf), Asn(Trt), Asp(O*t*Bu), Cys(Trt), Glu(O*t*Bu), Gln(Trt), His(Trt), Lys(Boc), Ser(*t*Bu), Thr(*t*Bu), Trp(Boc) and Tyr(*t*Bu), as well as Cys(S*t*Bu) for the thioesterification device.

Syntheses were performed at a 25 µmol scale. Protected amino acids (0.25 mmol, 10 equiv.) were coupled using HCTU (98 mg, 0.238 mmol, 9.5 equiv.) and *i*$Pr_2$NEt (87 µL, 0.5 mmol, 20 equiv.) in NMP (3 mL) for 30 min. Capping of potential unreacted amine groups was achieved by treatment with acetic anhydride (143 µL, 1.51 mmol, 60 equiv.), *i*$Pr_2$NEt (68 µL, 0.39 mmol, 15.5 equiv.) and HOBt (6 mg, 0.044 mmol, 1.8 equiv.) in NMP (3 mL) for 7 min. Fmoc group was removed by three successive treatments with 20% piperidine in NMP (3 mL) for 3 min. Deprotection and cleavage from the resin was performed through a treatment with TFA/$H_2O$/*i*$Pr_3$SiH/phenol (88:5:2:5) for 2 h, then precipitated by dilution into an ice-cold 1:1 diethyl ether/petroleum ether mixture, recovered by centrifugation, further washed three times with diethyl ether and dried under reduced pressure.

#### 3. Single fragment solid phase synthesis of the reduced form of BCR4

Sequence:

H-$^{1}$DFDPTEFKGPFPTIEICSKYCAVVCNYTSRPCYCVEAAKERDQWFPYCY$^{50}$D-OH

Synthesis of the reduced form of BCR4 was first attempted through a single fragment Fmoc SPPS, starting from Fmoc-Asp(O*t*Bu) TentaGel R PHB resin (132 mg, 0.19 mmol/g, 25 µmol). Tyr27 and Thr28 were introduced as a pseudoproline dipeptide (Fmoc-Tyr(tBu)-Thr(□$^{(Me,Me)Pro}$)-OH), and a double coupling procedure (2 x 30min) was used for residues Asn26 to Asp1.

Even using this optimized protocol, the target peptide was a minor component of the crude mixture, and we could not separate it from truncated acetylated peptides contaminants using standard semi-preparative HPLC.

**ESI-HRMS** (*m/z*): $[M+H]^+$ calcd. for $C_{266}H_{379}N_{62}O_{79}S_6$: 5897.5869, found: 5897.5911.

**HPLC analysis**: $t_R$ = 29.6 min (gradient: 5-50% B/A over 30 min).

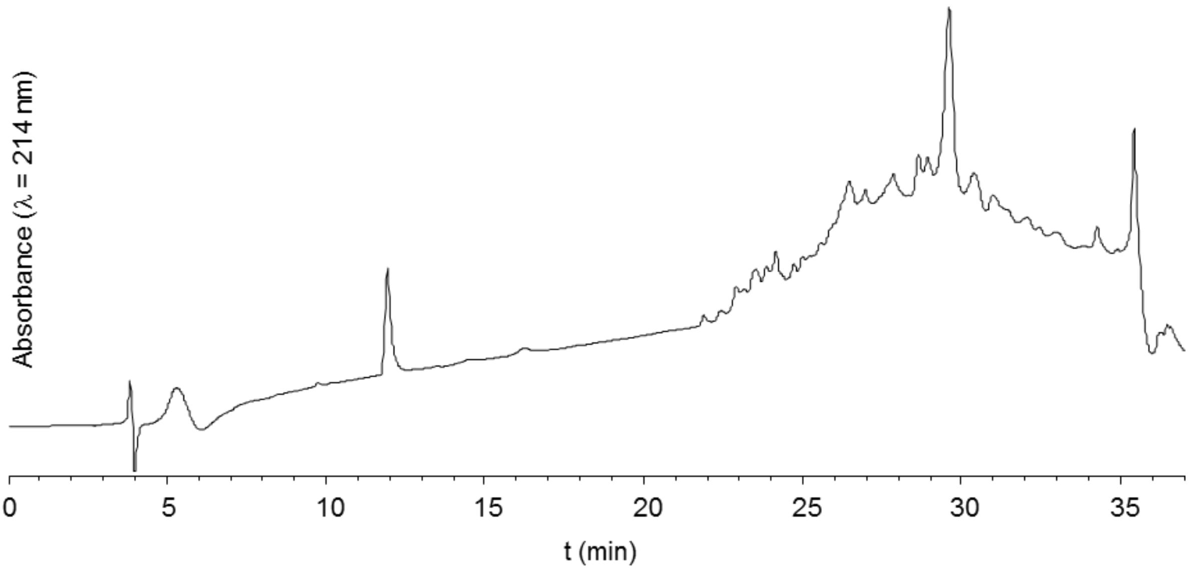


**Figure S1**: HPLC trace of crude reduced BCR4 obtained from a single fragment synthesis. Gradient: 5-50% B/A over 30 min.

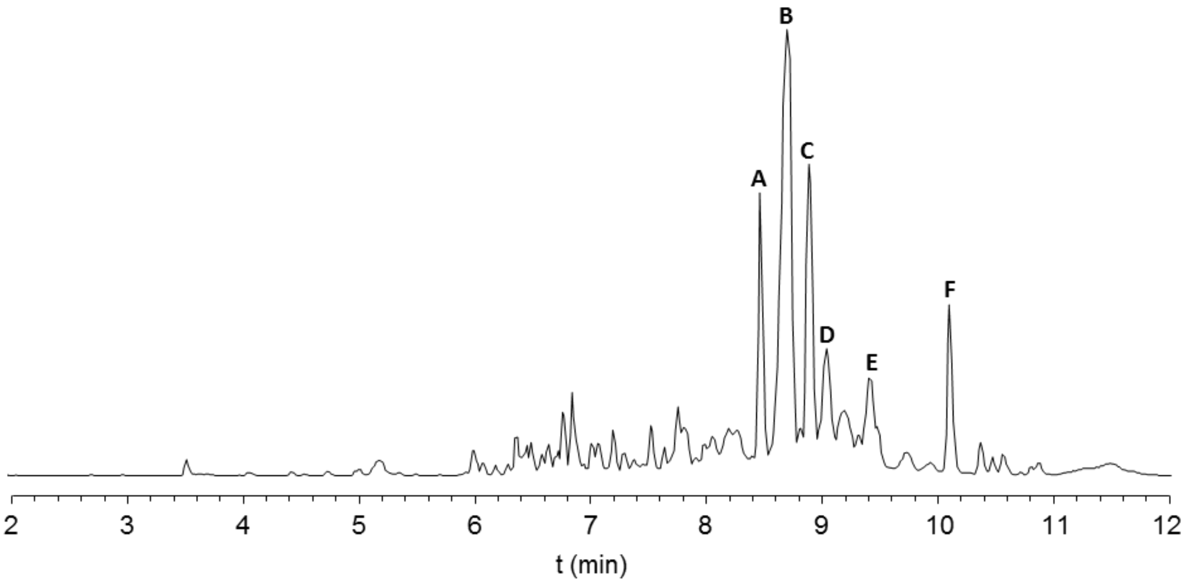


**Figure S2**: LC/MS analysis of crude reduced BCR4 obtained from a single fragment synthesis (base peak ion chromatogram).

**Table S1**: Attribution of the main peaks observed during LC/MS analysis of crude reduced BCR4 obtained from a single fragment synthesis.

| Peak ( $t_R$ (min) ) | $[M+H]^+$ calcd. | $[M+H]^+$ found | Attributed to |
|---|---|---|---|
| A (8.48) | 4561.9734 | 4561.9759 | Ac-[13-50] |
| B (8.71) | 4659.0262 | 4659.0286 | Ac-[12-50] |
| C (8.90) | 5897.5869 | 5897.5911 | [1-50]: reduced form of BCR4 |
| D (9.06) | 4715.0888 | 4713.0826 | Ac-[13-50] *t*Bu adduct |
| E (9.17) | 5953.6495 | 5953.6548 | [1-50] *t*Bu adduct |
| F (10.11) | 4742.0309 | 4742.0321 | Fmoc-[13-50] |

#### 4. Native chemical ligation-based synthesis of the reduced form of BCR4

Considering the difficulties observed during the single fragment SPPS, synthesis of the reduced form of BCR4 was achieved through a two-fragment native chemical ligation (NCL) strategy, based on the reaction of a [1-20] *N*-2-hydroxy-5-nitrobenzylcysteine (*N*-Hnb-Cys) crypto-thioester[1] with a [21-50] cysteinyl peptide (supplementary figure S3).

[1] (a) V. P. Terrier, H. Adihou, M. Arnould, A. F. Delmas, V. Aucagne, *Chem. Sci.*, **2016**, *7*, 339−345 (b) D. Lelièvre, V. P. Terrier, A. F. Delmas, V. Aucagne, *Org. Lett.*, **2016**, *18*, 920−923 (c) V. P. Terrier, A. F. Delmas, V. Aucagne, *Org. Biomol. Chem.*, **2017**, *15*, 316−319 (d) G. Martinez, J.-P. Hograindleur, S. Voisin, R. Abi Nahed, T. M. Abd El Aziz, J. Escoffier, J. Bessonnat, C.-M. Fovet, M. De Waard, S. Hennebicq, V. Aucagne, P. F. Ray, E. Schmitt, P. Bulet, C. Arnoult, *Mol. Hum. Rep.*, **2017**, *23*, 116−131.

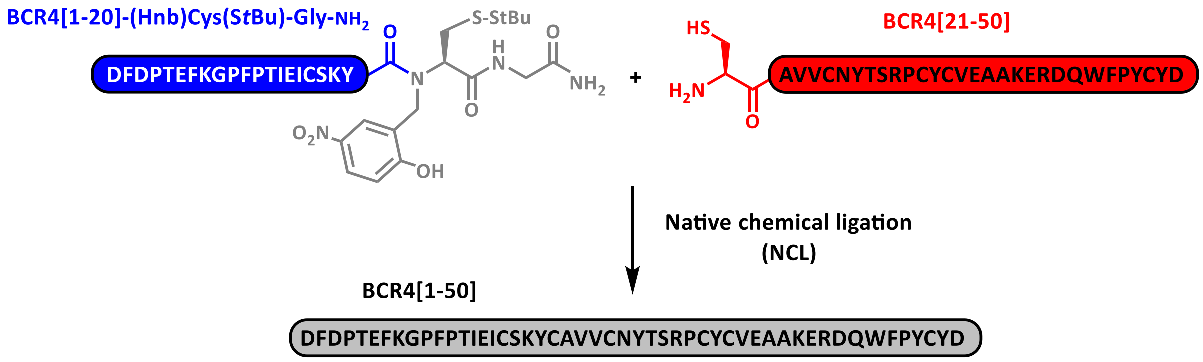


**<u>Figure S3</u>**: NCL-based synthesis of the reduced form of BCR4.

##### 4.1- Synthesis of BCR4[1-20] crypto thioester

Sequence: H-$^{1}$DFDPTEFKGPFPTIEICSK$^{20}$Y-(Hnb)C(StBu)G-$NH_2$

Rink linker, Fmoc-Gly-OH and Fmoc-Cys(S*t*Bu)-OH were successively coupled by automated SPPS on a Tentagel R $NH_2$ resin (120 mg, 0.21 mmol/g, 25 µmol). Resulting peptidyl resin was washed with a 1:1 DMF/MeOH mixture, then swollen in 9:9:2 DMF/MeOH/AcOH for 5 min. The reactor was drained off and the resin was washed with 1:1 DMF/MeOH. 2-Hydroxy-5-nitrobenzaldehyde (HNBA) in 1:1 DMF/MeOH (125 mM, 10 equiv., 2 mL) was then added and the reactor was left for 1 h under stirring through nitrogen bubbling. The reactor was drained and the resin was washed with 1:1 DMF/MeOH. Without delay, a fresh solution of sodium cyanoborohydride in 9:9:2 DMF/MeOH/AcOH (250 mM, 20 equiv. 2 mL) were added and the reactor was left for 1 h under stirring by nitrogen bubbling. The reactor was drained off and the resin was extensively washed with 1:1 DMF/MeOH, NMP, 20% piperidine in NMP, NMP, dichloromethane then NMP. Tyr20 was introduced through a 3 x 2 h coupling protocol, then the elongation from residues 19 to 1 was pursued using standard conditions, using a double coupling procedure for residues Pro12 and Phe11.

Crude peptide was purified by semi-preparative RP-HPLC to yield pure crypto thioester (15 mg, 4.9 µmol, 20%).

We found that the Asp3-Pro4 peptide bond was particularly sensitive to acid hydrolysis:[2] The crude or purified peptide should not be kept in HPLC solvents (0.1% TFA) for a prolonged time (>10 h) nor be heated above room temperature under these conditions, and the purified fractions should be lyophilized immediately after purification. Peptide re-dissolved in pure water was however stable for a few months at -20°C.

**ESI-HRMS** (*m/z*): $[M+H]^+$ calcd. for $C_{125}H_{179}N_{26}O_{37}S_3$: 2732.2087, found: 2732.2119.

**HPLC analysis**: $t_R$ = 19.6 min (gradient: 20-55% B/A over 21 min).

**HPLC purification**: $t_R$ = 17.4 min (Jupiter C4, gradient: 30-55% B/A over 19 min).

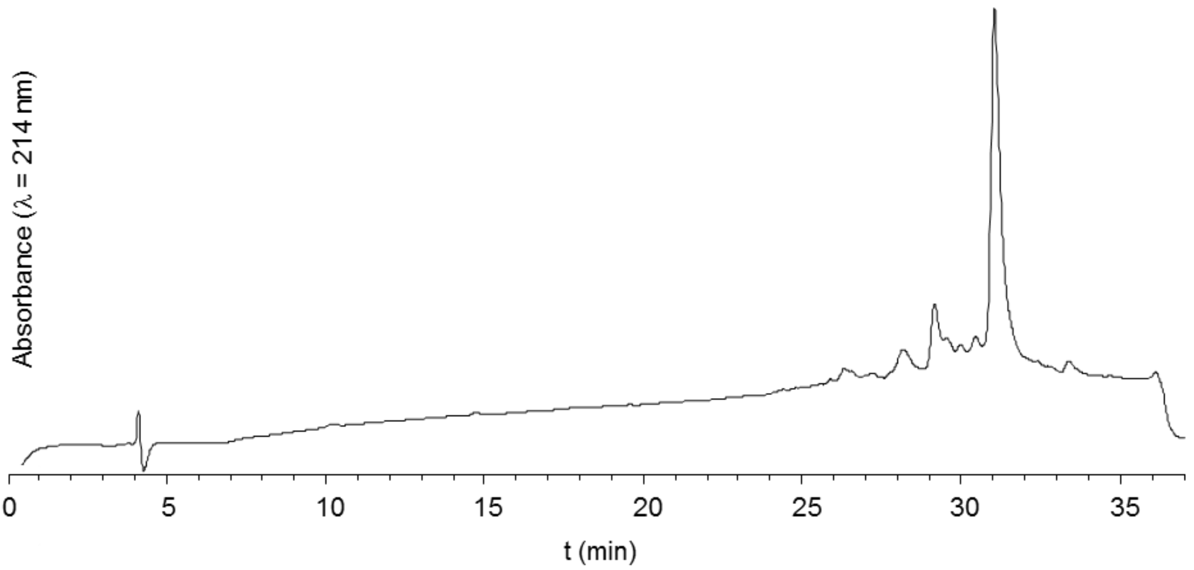


**Figure S4**: HPLC trace of crude BCR4[1-20] crypto thioester. Gradient: 5-50% B/A over 30 min.

[2] (a) D. Piszkiewicz, M. Landon, E. L. Smith, *Biochem. Biophys. Res. Commun.*, **1970**, *40*, 1173−1178 (b) I. Ségalas, R. Thai, R. Ménez, C. Vita, FEBS Lett., **1995**, *371*, 171−175.

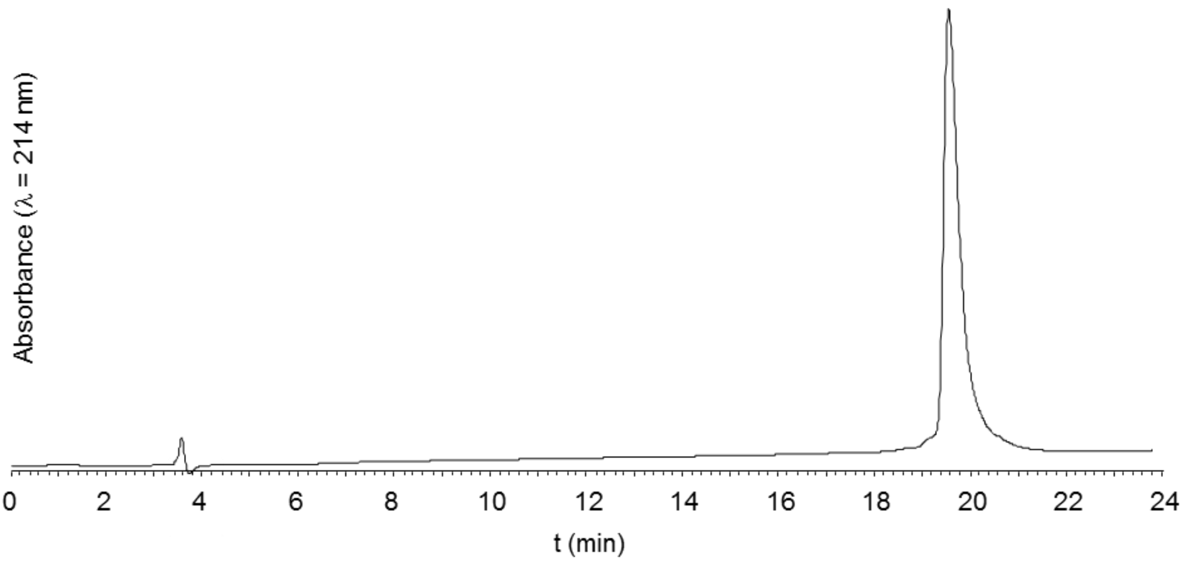


**Figure S5**: HPLC trace of purified BCR4[1-20] crypto thioester. Gradient: 20-55% B/A over 21 min.

**4.2- Synthesis of BCR4[21-50] cysteinyl peptide**

Sequence: H-$^{21}$CAVVCNYTSRPCYCVEAAKERDQWFPYCY$^{50}$D-OH

BCR4 [21-50] cysteinyl peptide was synthesized through standard Fmoc SPPS starting from Fmoc-Asp(O*t*Bu) TentaGel R PHB resin (132 mg, 0.19 mmol/g, 25 µmol), using a double coupling procedure for residues Ala39 to Val23.

Crude peptide was purified by semi-preparative RP-HPLC to yield pure cysteinyl peptide (16 mg, 4.0 µmol, 16%).

**ESI-HRMS** (*m/z*): $[M+H]^+$ calcd. for $C_{157}H_{225}N_{40}O_{47}S_5$: 3582.5049, found: 3582.5067.

**HPLC analysis**: $t_R$ = 13.9 min (gradient: 20-55% B/A over 21 min).

**HPLC purification**: $t_R$ = 14.2 min (Nucleosil C18, gradient: 25-30% B/A over 15 min, 50°C).

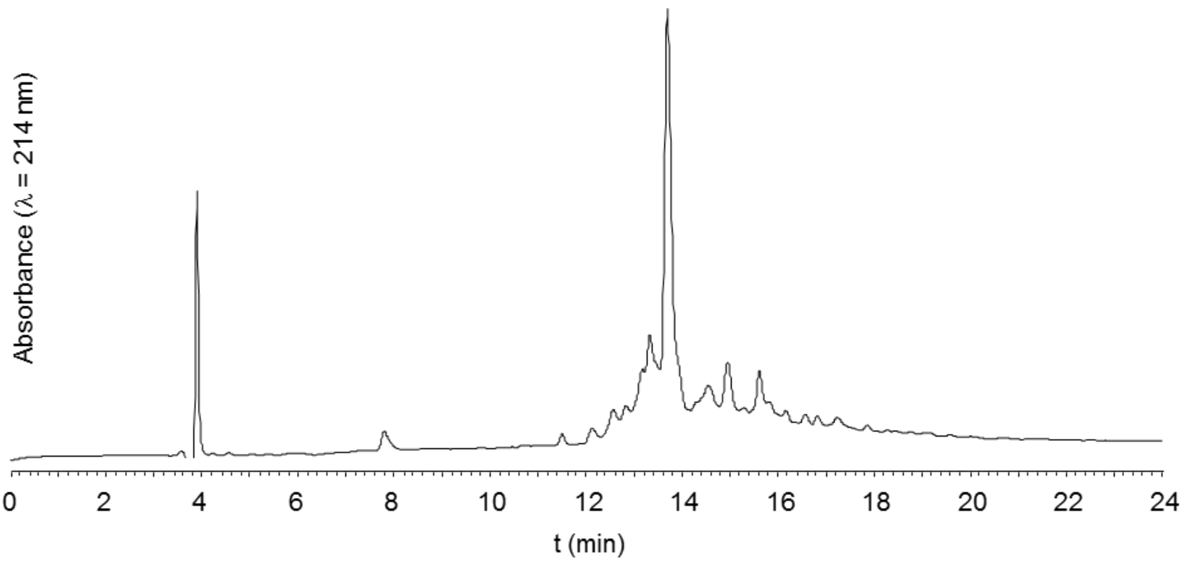


**Figure S6**: HPLC trace of crude BCR4[21-50]. Gradient: 20-55% B/A over 21 min.

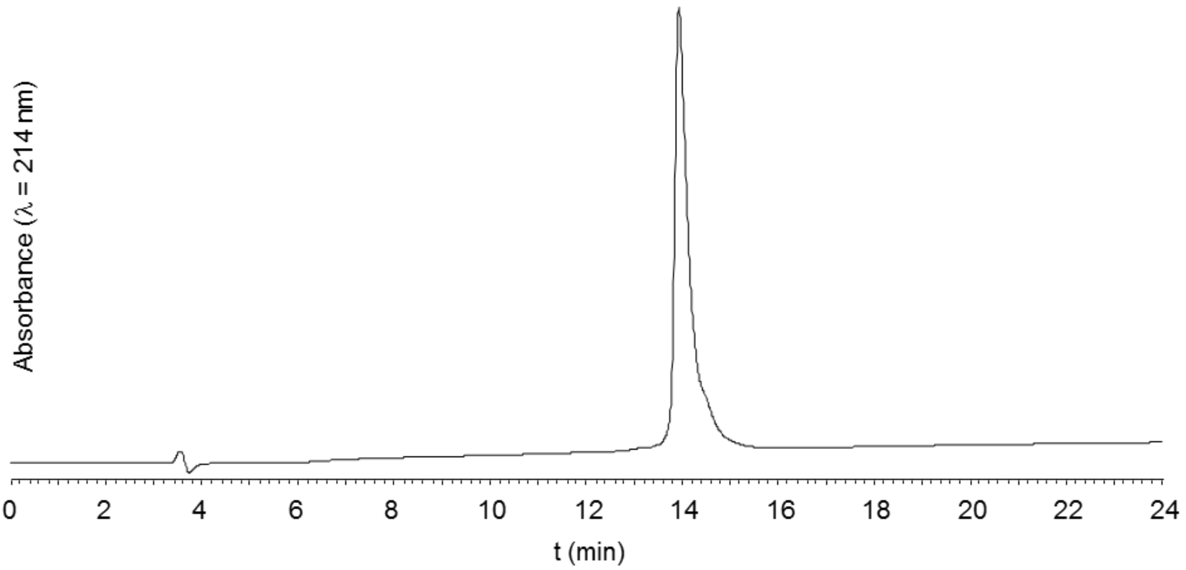


**Figure S7**: HPLC trace of purified BCR4[21-50]. Gradient: 20-55% B/A over 21 min.

##### 4.3- Native chemical ligation

Under an argon atmosphere, 765 µl of a deoxygenated 0.2 M sodium phosphate buffer pH = 6.5 containing 200 mM 4-mercaptophenylacetic acid (MPAA), 50 mM *tris*-carboxyethylphosphine (TCEP) and 6 M guanidine hydrochloride (Gu.HCl) was added to HPLC-purified [1-20] cysteinyl peptide (1.56 µmol, final concentration 2 mM), and 4.6 mg [1-20] crypto thioester (1.2 equiv.). The resulted solution was incubated at 37°C for 24 h, then quenched by addition of 15 mL of a $H_2O$/MeCN/AcOH 70:25:5 mixture. The solution was washed three times with 30 ml $Et_2O$ then centrifuged. The precipitate was dissolved in 1 mL 6M Gu.HCl, combined with the supernatant then purified by semi-preparative RP-HPLC to yield 3.3 mg (528 nmol, 34%) of pure BCR4.

The crude or purified peptide should not be kept in HPLC solvents (0.1% TFA) for a prolonged time (>10 h) nor be heated above room temperature under these conditions, and the purified fractions should be lyophilized immediately after purification. Peptide re-dissolved in pure water was however stable for a few weeks at -20°C.

**ESI-HRMS** (*m/z*): $[M+H]^+$ calcd. for $C_{266}H_{379}N_{62}O_{79}S_6$: 5897.5869, found: 5897.5883.

**HPLC analysis**: $t_R$ = 18.8 min (gradient: 20-55% B/A over 21 min).

**HPLC purification**: $t_R$ = 16.5 min (Jupiter C4, gradient: 30-53% B/A over 17 min).

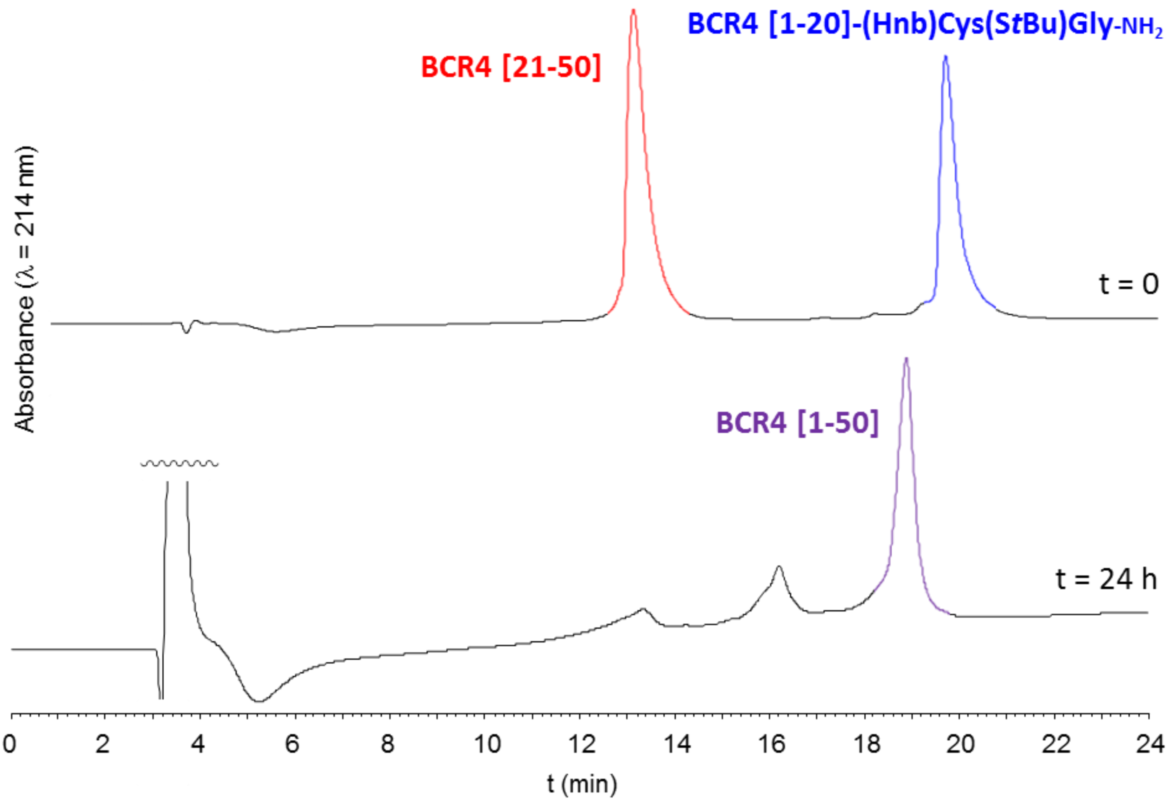


**Figure S8**: HPLC monitoring of the NCL reaction. Gradient: 20-55% B/A over 21 min.

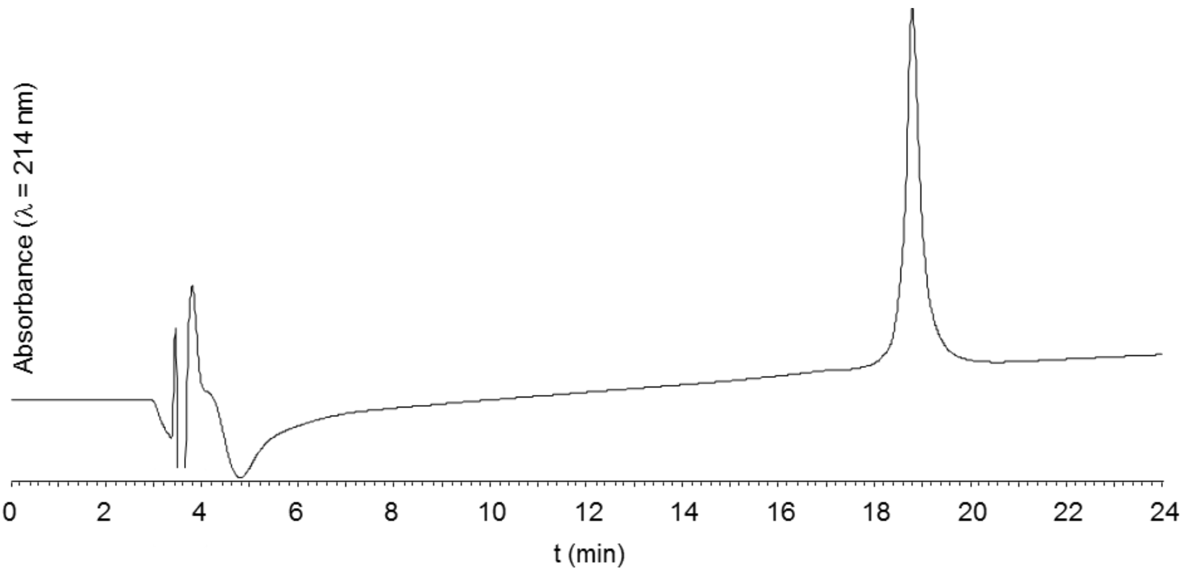


**Figure S9**: HPLC trace of purified reduced BCR4. Gradient: 20-55% B/A over 21 min.

#### 5. Oxidative folding

Oxidative folding was performed by incubating the reduced peptide (622 nmol) in 20.7 mL (30 µM final concentration) of a deoxygenated buffer containing 0.1 mM oxidized glutathione (10 equiv.), 1 mM glutathione (100 equiv.), 1 mM EDTA, 100 mM TRIS, pH 8.5, at 20 °C, for 48 h under an argon atmosphere. The reaction was acidified by adding TFA (200 µL), and the crude mixture was purified by semi-preparative HPLC to give pure BCR4 (135 nmol, 22%). The crude or purified peptide should not be kept in HPLC solvents (0.1% TFA) for a prolonged time (>10 h) nor be heated above room temperature under these conditions, and the purified fractions should be lyophilized immediately after purification. Peptide re-dissolved in pure water was however stable for several months at -20°C.

**ESI-HRMS** (*m/z*): $[M+H]^+$ calcd. for $C_{266}H_{373}N_{62}O_{79}S_6$: 5891.5400, found: 5891.5437.

**HPLC analysis**: $t_R$ = 22.5 min (gradient: 5-50% B/A over 30 min).

**HPLC purification**: $t_R$ = 25.7 min (Nucleosil C18, gradient: 5-45% B/A over 30 min).

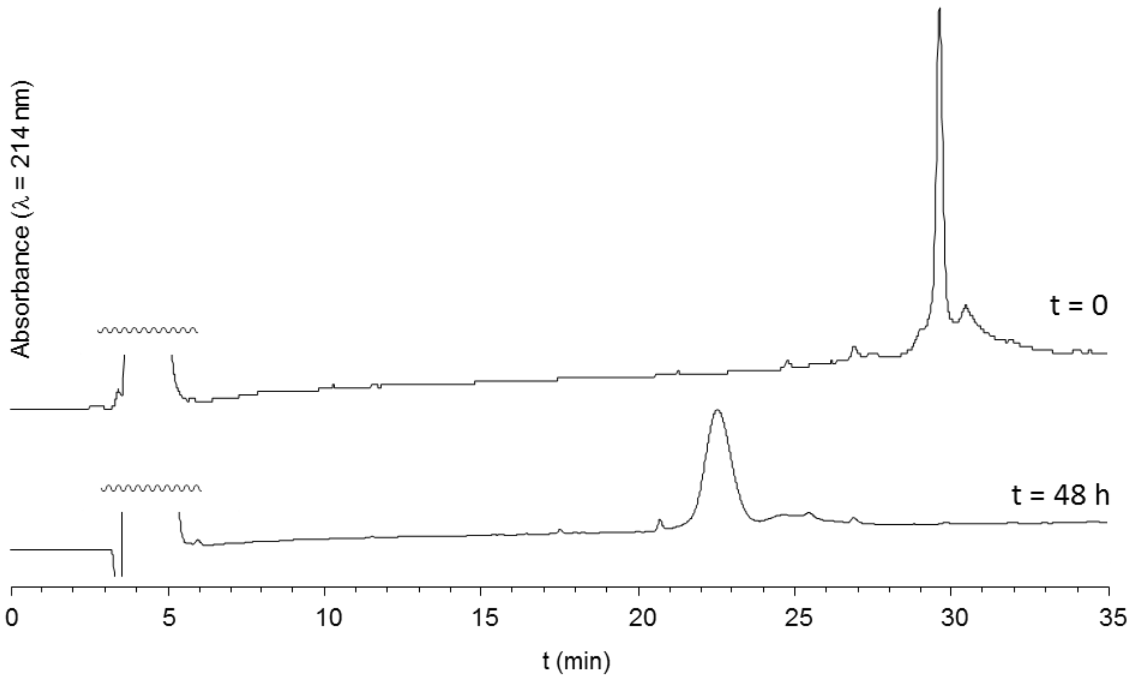


**Figure S10**: HPLC monitoring of the oxidative folding. Gradient: 5-50% B/A over 30 min.

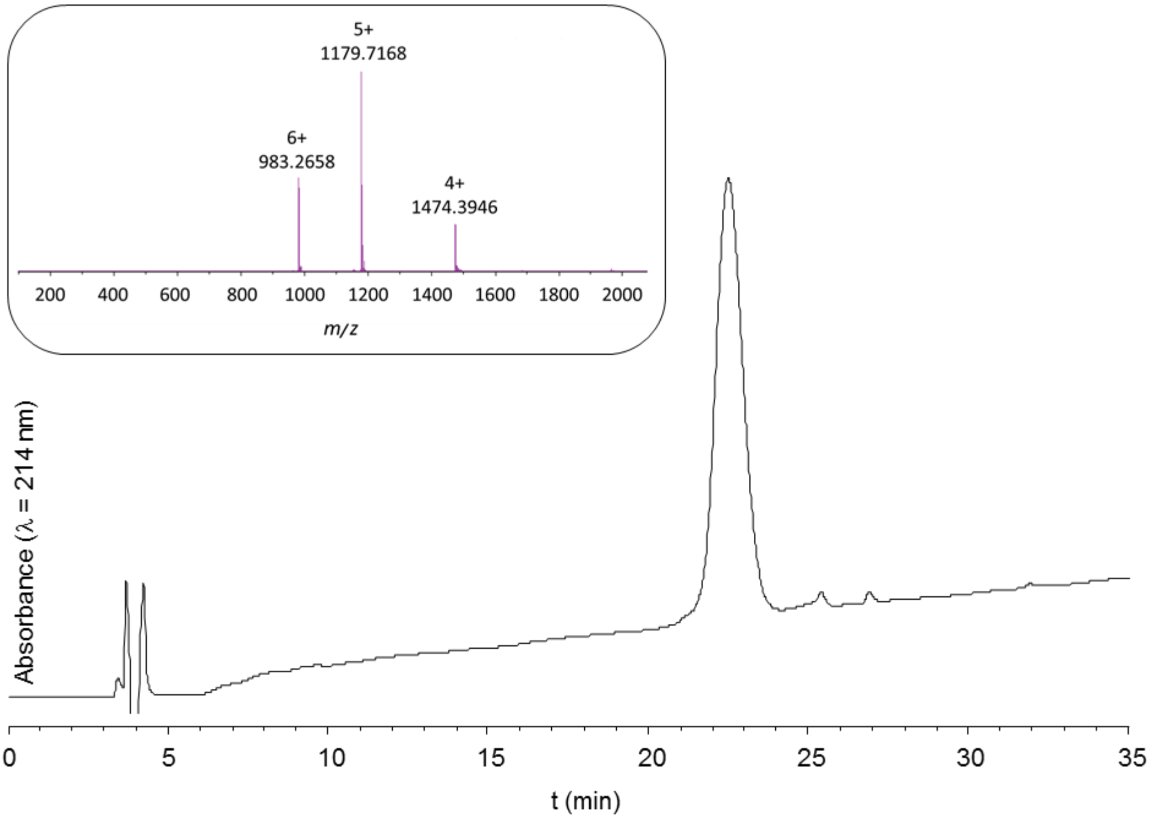


**Figure S11**: HPLC trace and ESI-HRMS spectrum of purified BCR4. Gradient: 5-50% B/A over 30 min.

**Table S2.** Protein sequences of the putative BCR homologs in 22 aphid species which sequences are deposited in publicly available databases.

| BCR subfamilies | Protein sequences in aphid species[a] |
|---|---|
| **BCR1-2-4-5** | **>Apis-BCR1**<br>MKLLHGFLIIMLTMHLSIQYAYGGPFLTKYLCDRVCHKLCGDEFVCSCIQYKSLKGLWFP<br>HCPTGKASVVLHNFLTSP<br>**>Apis-BCR2 (ACYPI38738)**<br>MKLLYGFLIIMLTIHLSVQYFESPFETKYNCDTHCNKLCGKIDHCSCIQYHSMEGLWFPH<br>CRTGSAAQMLHDFLSNP<br>**>Apis-BCR4**<br>MRLLYGFLIIMLTIYLSVQDFDPTEFKGPFPTIEICSKYCAVVCNYTSRPCYCVEAAKER<br>DQWFPYCYD<br>**>Apis-BCR5 (ACYPI084619)**<br>MRLLYGFLIIMLTIHLSVQDIDPNTLRGPYPTKEICSKYCEYNVVCGASLPCICVQDARQ<br>LDHWFAYCYDGGPEMLM<br>**>Apis-BCRnew1 NC_042494.1:96422603..96422806**<br>MRLLYGVLIIMLTIQLSVQYSYYPGRPFVSRHNCEAACTRICGFSNPCSCVQYGSIMWSP<br>HCRSGRAAGSWPGEDPY<br>**>Akon-FQ998496.1 BCR1 BCR2 BCR4 BCR5**<br>MRLLYVFLVVMLTMQLSIQYTSGSPFQTRYNCNNICHKLCGSAACACSQYRSLKGMWFPH<br>CANGQAAQVLHNFLSN<br>**>Akon-FO015834.1 BCR1 BCR2 BCR4 BCR5**<br>MKYFYGFLIIMLTIHLSVQYHYIESPFETRFGCDNVCYKLCGKRVPCSCVQYDAMNGLWF<br>PHCQEGHAAEELHQFL<br>**>Dnox-NW_015368581.1 BCR1 BCR2 BCR4 BCR5**<br>IRLLFGLLIIMLTVHLSIQEDDYPTRKQCNETCIANCRSDPNYEGRWMWCLREAGSEMIG<br>LWYCQC<br>**>Mros-WHPZ01509477.1 BCR1 BCR2 BCR4 BCR5**<br>MRFLYGFLIIMLTIHLSVQLSISPFEEKFTCDRICYKLCGNVNKCRCQQYDSLLNLWFPR<br>CSVGNAAIVLHEFLSNP<br>**>Mros-WHPZ01494279.1 BCR1 BCR2 BCR4 BCR5**<br>MRFLYGFLIIMLTIHLSIQLFISPFEEKFTCDRICYKLCGNVHKCRCQQYDSLLNLWFPR<br>CSGGNAAIVLHEFLSNP<br>**>Mros-WHPZ01338884.1 BCR1 BCR2 BCR4 BCR5**<br>MRFLYGFLIIMLTIHLSVQLSRSPLESRFECENICYSLCGGDNVCNCEQYKSLNNLWFPH<br>CRFGHAAMVLHEFLSSP<br>**>Mros-WHPZ01584390.1 BCR1 BCR2 BCR4 BCR5**<br>MRLMFGFLIIMLTINLSVQYSYYPGRPFISKYNCEAACTRICGFSNPCSCLQYDVISSLW<br>FPRCRSGHPAGV<br>**>Mros-WHPZ01569289.1 BCR1 BCR2 BCR4 BCR5**<br>MKLLFGFLIIMLTLHLSIQNPYHSDQSYRTKFECENDCSSMCITGYDGQCERLRTVYYLW<br>SCYCTPAG<br>**>Smis-CM017799.1-23116456 BCR1 BCR2 BCR4 BCR5**<br>MKLLFGFSIIMLTIHLSVQYSYYPGRPFASKYNCETVCTRICGFSNPCSCLQFDLMNAPL<br>WFPRCRSGHA |
| **BCR3** | **>Apis-BCR3 (ACYPI44142)**<br>MSVRKNVLPTMFVVLLIMSPVTPTSVFISAVCYSGCGSLALVCFVSNGITNGLDYFKSSA<br>PLSTSETSCGEAFDTCTDHCLANFKF<br>**>Agly-AG010439-PA BCR3**<br>MWTFILVGLLMMTCVTEASRLNRFMSNVCYFGCLAKRVACFSSTGAIFGTVPYGIIAVTP<br>ALESCTVVFRICKASCIAILILPKI<br>**>Dnox-NW_015368357.1 BCR3**<br>MQSRSNVVWPSLFVALLMMSPVTHANIFIAAVCYSGCGSLALVCYVSNGIANGIHYAKTF<br>TSLPPTEVGCGEAFVECKNNCLEHF<br>**>Dnox-XP 015363544.1 PREDICTED:**<br>**uncharacterized protein LOC107161588] BCR3** |

218

| | |
|---|---|
| | MVAQKMWSLILIGLLFSSSANAGPIXAGICYAGCAAVTVACFGAAGFTFGTVPGAVIAAT<br>PALAACNAAFGICEASCVAALVVPTP<br>**>Mper-MYZPE13164_0_v1.0_000159160.2_pep \| 88 aa BCR3**<br>MSARNNMLPSLFIVLLIMSPVTPAHFFIAYVCYAGCGSLAIVCYVSNGIANGIQFAKTAS<br>ALPPTKISCGQAFNACKDDCLVNFTLTP<br>**>Mcer-Mca00644.t1-protein BCR3**<br>MSARTNVLPSLFVMLLIMSPVAPAHLFIAYVCYAGCGSLALTCYVSNGLANGLEFIKTAS<br>FLPPTAITCAEAFVACKTDCLENFTLTP<br>**>Tcit-CD450908.1 BCR3**<br>MSARKNILATVFIVLLFSTSVTPTNIFISAVCFSGCGTMLLTCYVITGFEFIKAAVFDDH<br>LPTSGCRTMYHECTAVCKKSYK<br>**>Rpad-Rp1008 BCR3**<br>MSAQNKMLTSLFIVLLITTSVTPTSIFISSICYSGCGTMAVACYVMTGIQYIMGASGLTT<br>NGMGCGASFVVCKDSCIESYN<br>**>Rmai-NC_040880.1 BCR3**<br>MSAQNKMLTSLFIVLLITTSVTPTSIFISSICYSGCGTMAVACYVMTGIQYIMGASGLTR<br>TGLGCGESFVVCKDSCIQSY<br>**>Rmai-XP_026822757.1 BCR3 uncharacterized protein**<br>**LOC113560848**<br>MVGQKMLSLLMVGLLLSPPALAGPVAAGICYAGCAAVTVACFTAAGFTFGTVPASVIAAT<br>PVLAACNTAFGVCEASCVAALVVPTP<br>**>Rmai-NC_040879.1 BCR3**<br>MWTFMMAGLLMMTCVTEAGPFNRLISNVCYLGCLAERVACFSSSGAIFGTVPYGIIAVTP<br>ALNTCNAIFKICKVGCIAILISPR<br>**>Sgra-QEWZ01000258.1 BCR3**<br>MSAQNKMLTSLFIVLLMTTSVTPSSIFISSICYSGCGTMAVACYVMTGIQYIMGASGLTR<br>IGLSCGESFFVCKDSCIQSY<br>**>Sgra-QEWZ01001431.1 BCR3**<br>MWTFIMAGLLMMTCVTEAGPLNRLMSNVCYLGCLAERVACFSSSGAIIGTVPYGIIAVTP<br>ALNTCTAIFNICKVGCIAILISPR<br>**>Msac-NW 020270519.1 BCR3**<br>MSARNNIFTSLFVVLLMTTSVTPAKIGLSIICYSGCGSVAIGCYVINSFEFLKSFFVPDT<br>MTSDCGESFDECKYACI<br>**>Msac-XP 025200550.1 BCR3 uncharacterized protein**<br>**LOC112598338**<br>MVAQKMLSLVLVGLLLSSSAHAGPLTAGICYAGCAAVTVACFSAAGFTFGTVPGAVIAAT<br>PALAACNAAFGICEASCVAALVVPVP<br>**>Msac-XP 025194587.1 BCR3 uncharacterized protein**<br>**LOC112594148**<br>MVAQKMLSLVLVGLLLSSSAHAGPLTAGICYAGCAAVTVACFSAAGFTFGTVPGAVIAAT<br>PALAACNAAFGICEASCVAALVVPVP<br>**>Agos-DR392815.1 BCR3**<br>MSAKKNILATAFIVLLFSTSVTPTNIFISAVCFSGCGTMLVTCYVITGLEILKAAVVDDH<br>LPTSGCRSMFHECKDVCYRSYR<br>**>Agos-XP_027838196.1 BCR3 uncharacterized protein**<br>**LOC114120478**<br>MVAHKMLSLVLVGLLLSSTAHAGPLAAGVCYAGCAAVTVACFSAAGFTFGTVPGAIIAAT<br>PALAACNAAFGICEASCIAALVVPVP<br>**>Acra-KAF0749826.1 BCR3**<br>MTAQKMWTFVLVGLLMMTCVTEASRLNRFMSNVCYFGCLAERVACFSSTGAIFGTVPYGI<br>IAVTPTLESCTVVFKICKASCIAILISSKI<br>**>Acra-KAF0768549.1 BCR3**<br>MVTHKMLSLVLVGLLLSSSAHAGPLAAGVCYAGCAAVTVACFSAAGFTFGTVPGAIIAAT<br>PALAACNAAFGVCEASCIAALVVPVP<br>**>Acra-VUJU01000019.1 BCR3**<br>MTAKKNILATVFIVLLFSTSVTPTNIFISAVCFSGCGTMLVTCYVITGLEIIKAAVVDDH<br>LPKSGCKAMFQECKAICVKSYK<br>**>Cced-VVC36620.1 BCR3**<br>MFARKTIVLLTVLMMFGIAQAGPLAAGICYAGCASVTVACFAAAGFTFGTVPGAVIAATP<br>ALAACNTAFGICEAACVAALVAPTV<br>**>Pnig-scaffold 705 BCR3** |

| | |
|---|---|
| | MSARKDVLLSVFVAMLMMSSVAPTHILVSVVCYSGCGSMALVCYLSTGIANGLHPSKSTE<br>SSNADCGKSYDSCISDCVLNF<br>**>Pnig-scaffold_2336 BCR3**<br>MLSLILVGLLMMANATEAGRLASNFCYFGCLAERVACFSSSGAILGTVPFGMIAVAPALK<br>TCTAIFGVCKASCFALLLMPII<br>**>Mros-WHPZ01586055.1 BCR3**<br>MSARKNVLPILFVVLLIMSPVTPTSIFISAVCYSGCGSLALVCYVSNGVTIGMDYIKSTV<br>TLPSELTCGEAFDECKDNCLSTFSF<br>**>Mros-WHPZ01559067.1 BCR3**<br>MLSLMLVGLLIASSANAGPLAAGICYAGCAGVTVACFSAAGFTFGTVPGALIAATPALAA<br>CNAAFGICEASCMAALFVP<br>**>Mros-WHPZ01582937.1 BCR3**<br>IWSLMLAGLLIASSANAGPIAAGICYAGCAAVTVACFAAAGFTFGTVPGAVIAATPALAA<br>CNAAFGICEASCVAALVVP<br>**>Asol-PVMI01067726.1 BCR3**<br>MWVKNNMLPSLFIVLLIMSPVTPTSFFIAYVCYAGCGSLALVCYVSNGISNGIQYVKTASV<br>LPPAEIGCAEAFGDCKTDCLENF<br>**>Asol-PVMI01040116.1 BCR3**<br>MWSLILVGLLISASANAGPIAAGVCYAGCAAVTVACFSAAGFTFGTVPGAVIAATPALAAC<br>NAAFGVCEASCMAALFVP<br>**>Smis-CM017802.1 BCR3**<br>MSAQKNVLPALFIMLLIMTPVTPTNVLISAVCYSGCGSLALVCYVYNGISIGVDYFKSSDP<br>LPSLESSCGRSFKRCKDHCLKTFTF<br>**>Smis-SSSL01000270.1 BCR3**<br>LWSLIFVGLSISSSANAGPIAAGICYAGCAAVTVACFSAAGFTFGTVPGALIAATPVVAAC<br>NAAFGICEASFITALIVP |
| **BCR6** | **>Apis-BCR6 (ACYPI49532)**<br>MDLFKKFCFVYLILHLTLLLFVDSSDYDDYEERKKYNGSVPNENKTCLIAWETSIMSEPT<br>PTCWIMCKIRCILLSRTTQWRCKISNNQIWENCHCCNDDTSYATFDFY<br>**>Dnox-NW 015369358.1 BCR6**<br>MNLLKKAYFVYFIFIFSLLYGDSYESDEKRAKYNDEDKNSEKTCNVPWSTSVFPEPMTSC<br>YFFCKKRCQALSFTSQWRCEEIVSFAITKCQCCKGEFSNYIYNMY<br>**>Dnox-XP_015376899.1 PREDICTED: uncharacterized protein LOC107171180 [Diuraphis noxia] BCR6**<br>MNLLKKACFVYFILVLSLLFVDSFEDGEKRAKYNGDAPNDNRTCNIPWKTSLFSEPYSSC<br>WVMCKGRCIFLSRTTQWRCKKSKYDLLGNCHCCNDDTLNVYFDL<br>**>Mper-MYZPE13164_0_v1.0_000200120.1_pep \| 104 aa BCR6**<br>MSLLKKACFVYFIFTLTLLFVSSYEDYERRAKYNADDENDNKTCNIPWETSLMSEPYPSC<br>WLMCKGRCIILSRTTQWRCKMSPNEIFGNCHCCCDDNVNVIYDF<br>**>Mcer-Mc581 BCR6**<br>MNLLKKACFVFFILTLTLLLVSSYEDNEKRAKYNGDDANDNKTCNVPWETSLMSEPYPSC<br>WLMCKGRCILLSRTTQWRCKMSPNEIFGNCHCCGGE<br>**>Save-JK721916.1 BCR6**<br>MSLLRKFCIICLILNLTFLLFADSYDDVDYEPLKKYDGDVPNDNKTCGVTWRTSWLSELT<br>PSCWIVCKVRCIILHRTTQWRCKKSDNPMWENCHCCTD<br>**>Masc-FO024986 BCR6**<br>MNPLKITCFIYFIVLLMSFCVYSKEEVYSKYYDDVPNDNKTCYFPWKTSILPEPYPTCWL<br>MCRLRCIVMSRTSQWRCKKSNHNLQGNCNCCTDN<br>**>Pnig-scaffold 992 BCR6**<br>VYFIYFIVLLMSVGLDSEEEENFSKYYGDMSNDNKTCYDPWQKSLLSEPFLVCWSSCKIR<br>CFILSGTGQFRCKKSYFGVVGNCQCCRDN<br>**>Pnig-scaffold_4247 BCR6**<br>MYLINKTSFISFILLSLICVHVNSDCRGAYNDSATDDKKCCPVAWTTPQLTVIKESYTQC<br>ANNCKTRCLNKRKTPQWQCLANTFTTLYSNCRCCTGEIRKLTYK<br>**>Pnig-scaffold 6689 BCR6**<br>MYLINNTSFIFFILLSLVCVNDAENDTCVYKDSDPNNKKCCPNMGWTNVCLDSSGKPKAI<br>DYSNCISNCQSDCNNKKGTKEWACAQMGSLYKCMCCVSEILNKL<br>**>Pnig-scaffold_18197 BCR6**<br>MYFINKTSFISFILLSLVYMHVNGDCLSVDECCSYNWTALLITTPFECDHGCFSNCQSIE<br>HTQNWYCVQDHRHNTGTCYCCTGEIHKLTY |

| | |
|---|---|
| | **>Pnig-scaffold_4321 BCR6**<br>MYLINNTSFIFFILLSLICVNDATNDCCVYNESAPNEKRCCPNDWKEPRLDAAGKYISQE<br>YSTCLCTCKAECKHYLNIQKWACTPDMDYHVCMCCVSEILN<br><br>**>Mros-WHPZ01589938.1 BCR6**<br>MNLLKKICFIYLILNLTFFLFVDSYYDDYEERKKYDGNLPNDNKTCEIPWETSIMSEPTP<br>TCYIMCKVRCIILSRTVQWRCKASSNGIWENCHCCNGE<br>**>Mros-WHPZ01474234.1 BCR6**<br>MYFINNLGLFFLILFTLAYVNCDEKGPYSSHDDSEHKKCKIDWVRATDGNGHIMSCSVKC<br>QTKCRYYQNTDQWRCRKSSTGLTKTCECCRGE<br>**>Mros-WHPZ01235886.1 BCR6**<br>MYLINNLSLFFLILFTLAYVNCDDERGPYHSSAEDDQKQCMVKWVKATHGGGNIASCLFC<br>KLKCKKKKKTSQWRCRKSKSGLTKKCECCTGE<br>**>Asol-PVMI01043125.1 BCR6**<br>MGLLKKACFVYFILTLTLLFVSSYEDYEKRAKYNDYDENENKTCNVPWETSMMSEPYSSC<br>WLMCKGRCIILSRTSQWRCKKSHHEILGNCHCCGGE<br>**>Smis-CM017798.1 BCR6**<br>MSLLRKFCIICLILNLTFLLFADSYDDVDYEPLKKYDGDVPNDNKTCGVTWRTSWLSELT<br>PSCWIVCKVRCIILHRTTQWRCKKSDNPMWENCHCCTGE<br>**>Smis-CM017799.1-7773692 BCR6**<br>MFLINNSGLIFLILFTLAYVNCDDTPGPYNSGDESDNKKCTIPWVPVTPEEGDTSTCLL<br>KCQTKCSDSQQTDQWRCKSLKKCECCRGE |
| **BCR8** | **>Apis-BCR8**<br>MSGYAKLLIFAFLLVLSVSQVLGCRGQCWKDVKPRDDFCSEIFRYQYTTMAPANVLCYCC<br>RRFIVED<br>**>Agly-AG001416-PA \| 80 aa BCR8**<br>MYRYTKVVVFVFILTLSANLANSSSMTTEGYKCPRSHCWTEKEPRDEFCSTIFRYEFATI<br>ELANVFCYCCRRLGSFILQ<br>**>Dnox-XR_001505997.1 BCR8**<br>MSHNMKLVIFAFLLILSVCQACGCRNNCWTDIKYRDDYCSELFRYQYTTMDPANVLCYCC<br>RRL<br>**>Mper-MYZPE13164 0 v1.0 000003380.1 pep \| 66 aa BCR8**<br>MNRNVKLVIFAFLLILSVSQVLGFGCPRGQCWIDKKKRDDFCLEIFRYEHTTMDPANVLC<br>FCCRRL<br>**>Mcer-Mc1616 BCR8**<br>MNHNVKLLIFAFLLILSVSQVLGCRGQCWIDIKPRDDFCSEIFRYQYTTIAPENVLCYCC<br>**>Akon-FQ999398.1 BCR8**<br>MNGHAKLLIFAFLLILSVSQVLGCRGQCWEEVKPRDDFCSEIFRYQYTTMEPANVLCYCC<br>RRFKLE<br>**>Masc-FO018431.1 BCR8**<br>MNRFTQLLIFAILLVLTISQVSACRGNCWTDEKYRDSYCSEIFRYKYRTFDVANVMCHCC<br>RSVI<br>**>Rpad-FO059758.1 BCR8**<br>MNRYIQLLVFVILLTLSISQVSGCRGQCWTDVKFRGEFCSQIFRYVYTTMEPANVVCYCC<br>RR<br>**>Rmai-NC_040878.1 BCR8**<br>MNRYLQLLVFVILLTLSVSQKSGCRGQCWTDVKFRDEFCSEIFRYVYTTIEPANVVCYCC<br>RR<br>**>Agos-NW 021007069.1 BCR8**<br>MYRYTQLVVFVFILTLSASLANSSSVTTEGYKCPRRQCWTEVEPRDEFCSEIFRYEFTTK<br>EPTNVFCYCCRR<br>**>Msac-NW_020271346.1 BCR8**<br>MNRYTQLLVLVFLLTLTISQVLGKCRDECWIDFRIRDDSCPLLFRYQYVTAAPANILCFC<br>CR<br>**>Acra-KAF0753610.1 BCR8**<br>MYRYTQLVVFVFLFTLSVSLSKSISVTTEGYKCQRGQCWTEVEPRDDFCSEMFRYEFITL<br>PPANVLCYCCRRI<br>**>Acra-VUJU01015826.1 BCR8**<br>MYRYMRLVVFVFLLTLSVILARSAPMAEGDTCYRGQCWTEVKPRDDFCSEIFRYDFTSKA<br>NVLCYCFRR |

| | |
|---|---|
| | **>Acra-VUJU01005956.1 BCR8**<br>MYRYMQLVVFVFLLTLSVSLAKSDPIVEGDTCFRGQCWTEVKPRDDFCTDIFRYNFTSKA<br>NELCYCCRR<br>**>Pnig-scaffold 94 BCR8**<br>MNRYTQLLIFAILLILTVNQALACRGNCWIDEKYRDSFCSEIFRYKYKIPNPVNILCHCCRR<br>**>Mros-WHPZ01587143.1 BCR8**<br>MNGQAKLLIFAFLLILTVSQVLGCRGRCWEDVKFRDDFCSEIFRYQYTTMKPAKALCYCCRRFKIE<br>**>Mros-WHPZ01588751.1 BCR8**<br>MNRTTYMVILAIIVVFCLSVTVMGCETNCWLNDWTTRDSACNGRVRYSYPGPSNGRCYCCQ<br>**>Asol-PVMI01009763.1 BCR8**<br>MNCNVKLLIFAFLLILSVSHVLGCRGNCWIDLKYRDNFCSEIFRYQYTTMEPANVLCYCCRR<br>**>Smis-CM017797.1-29839389 BCR8**<br>MNSTVKLLIFAFLLILSVSQVLGCKGQCWKDAEPRDDFCSQEFRYQYLTSKPANVLCYCC<br>**>Smis-CM017797.1-29874451 BCR8**<br>MNRTTYMVILAIIVIVCLSVTVMGCEKNCWLNDWRTRDAACNDRVKYSYPGPVHGKCYCCR |

[a] **Abbreviations:** Acra, *Aphis craccivora*; Agly, *Aphis glycines*; Agos, *Aphis gossypii*; Akon, *Acyrthosiphon kondoi*; Apis, *Acyrthosiphon pisum*; Asol, *Aulacorthum solani*; Cced, *Cinara cedri*; Dnox, *Diuraphis noxia*; Masc, *Myzus ascalonicus*; Mcer, *Myzus cerasi*; Mper, *Myzus persicae*; Msac, *Melanaphis sacchari*; Mros, *Macrosiphum rosae*; Pnig, *Pentalonia nigronervosa*; Rmai, *Rhopalosiphum maidis*; Rpad, *Rhopalosiphum padi*; Save, *Sitobion avenae*; Sgra, *Schizaphis graminum;* Smis, *Sitobion miscanthi;* Tcit, *Toxoptera citricida*. No BCR were found in Elan, *Eriosoma lanigerum* and *Sfla, Sipha flava.*
